# *C. elegans* display antipathy behavior towards food after contemporaneous integration of nutritional needs and dietary lipid availability

**DOI:** 10.1101/2024.02.23.581740

**Authors:** Nicole L. Stuhr, Carmen M. Ramos, Chris D. Turner, Alexander A. Soukas, Sean P. Curran

**Affiliations:** Leonard Davis School of Gerontology, University of Southern California, 3715 McClintock Ave, Los Angeles, CA, 90089 USA; Dornsife College of Letters, Arts, and Science, Department of Molecular and Computational Biology, University of Southern California, 1050 Childs Way, Los Angeles, CA, 90089 USA; Center for Genomic Medicine, Massachusetts General Hospital, Boston, MA 02114 USA

**Author notes:** Corresponding author: Sean P. Curran.

**Keywords:** diet, behavior, AWC, AWB, neurons, aging, lipid, vaccenic acid, sphingosine rheostat, *C. elegans*, *Methylobacterium*

## Abstract

Organisms utilize sophisticated neurocircuitry to select optimal food sources within their environment. *Methylobacterium* is a lifespan-promoting bacterial diet for *C. elegans* that drives faster development and longevity, however after ingestion, *C. elegans* consistently choose any other food option available. A screen for genetic regulators of the avoidance behavior toward *Methylobacterium* identified the AWB and AWC sensory neurons and the *odr-1* guanylate cyclase expressed exclusively in those four ciliated neurons as mediators of the antipathy response. Metabolic profiling of the *Methylobacterium* diet reveals a macromolecular profile enriched in saturated fats and here we show that *C. elegans* sense and integrate signals related to the type of ingested lipids that subsequently cues food-related behaviors. Moreover, disruption of endogenous lipid metabolism modifies the intensity of antipathy toward *Methylobacterium* which suggests that the current state of lipid homeostasis influences food preference. Enhanced expression of the sphingolipid degradation enzyme Saposin/*spp-9* enhances antipathy behaviors and activation of the sphingosine rheostat and more specifically modulation of the bioactive lipid mediator sphingosine-1-phosphate (S1P) acts as a signal to promote avoidance of *Methylobacterium*. Taken together, our work reveals that *C. elegans* modify food choices contemporaneously based on the availability of dietary lipids and the ability to metabolize dietary lipids.

**HIGHLIGHTS:**

- Uncover new molecular mechanisms underlying the decision matrix an animal uses to choose what foods to eat.
- Define the molecular mechanisms underlying an antipathy behavioral response toward foods after initial ingestion that contemporaneously integrates dietary needs with nutritional profile.
- ODR-1 signaling from AWB and AWC ciliated neurons of the *C. elegans* nervous system mediate the antipathy response to diet.
- Manipulation of sphingosine-1-phosphate (S1P) of the sphingosine rheostat controls the intensity of the antipathy behavioral response.
- Modulating antipathy behaviors can impact the magnitude of the lifespan-promoting effects of longevity diets.

## INTRODUCTION

Food is imperative to fuel growth and essential cellular functions for all organisms but not all foods are created equal [1]. The relationship that an individual has with food is complex and requires balancing taste and nutritional needs but is always bounded by the limitations of currently available sources. Throughout life, an individual’s nutrition can change due to organismal needs, the evolution of food preference, food availability, and age-related changes in utilization. Organisms must learn to distinguish between diets of different quality to have the best chance for survival[2–4]. In the short term, food choices can impact cellular metabolism while over longer periods can influence aging and age-related diseases, which has led to a substantive focus on elucidating how diet exerts such profound influence over health and longevity [5].

*C. elegans* inhabit a wide range of environments that span diverse climates and terrains[6, 7] but are most often found in habitats rich in microbial species. In nature, *C. elegans* can feed on many species of bacteria found in the soil, ranging from *Proteobacteria, Firmicutes, Bacteroidetes,* and *Actinobacteria* while in the laboratory, *C. elegans* husbandry typically utilizes a monoculture of *Escherichia coli* [7, 8]. Even with the lack of bacterial diversity in *C. elegans* laboratory environment, previous studies have shown that altering the *E. coli* species used for experiments can impact physiology in the worms, primarily linked to the different nutrient composition of each diet [9–12]. Bacterial diets supply macronutrients to the worms, all of which are needed to generate energy for cellular processes. Gross analysis of these macronutrients [13, 14] and metabolites [11, 13] have identified distinct differences in the makeup of the food sources, which likely contribute to the differing physiological attributes associated with each diet including, and not limited to, lifespan [15–17], healthspan [18–20], and reproduction [21–23].

The influence of diet on life history traits like development, reproduction, survival, and aging are facilitated in part by genetics [24–26]. The discovery of “gene-diet” pairs where the function of a gene becomes discernable only on a particular diet[24] has sparked excitement to define how an individual’s genetic profile can be leveraged to inform diet selection as a component of precision health and aging [27, 28]. Despite recent advances, the molecular mechanisms underlying diet choice and utilization require additional study.

Across organisms, the host microbiome plays critical roles in metabolism and health [29–31]. In addition, exposure to pathogenic bacteria can negatively impact health and like most organisms, *C. elegans* can learn from previous experiences and exposure ways to detect and avoid future interactions with potential pathogens [32–34]. Host-microbe interactions are complex and behavioral responses are regulated by distinct neurocircuitry [35, 36]. Our understanding of how an organism can distinguish between beneficial and detrimental food sources suggests selection for diets that promote fitness advantages [37, 38]. However, in this study we reveal a negative association, and behaviors that avoid *Methylobacterium,* which is not pathogenic, but also has been documented to promote health and longevity. We identify ODR-1 signaling in the AWB and AWC sensory neurons as the mediator for negative preference of the *Methylobacterium* diet and demonstrate that the antipathy for this food is associated with context dependent associations between the lipid composition of the diet and the current state of intracellular lipid homeostasis that is relayed in part by sphingosine-1-phosphate signaling. Ultimately, our work reveals a better understanding of the molecular basis underlying how an organism perceived the composition of a food and the impact it can have on dietary decisions.

## RESULTS

### *C. elegans* exhibit avoidance behavior away from longevity-promoting *Methylobacterium*

*C. elegans* must adapt to a variety of microbes in their natural environment and distinguish between what food sources are beneficial and detrimental[2, 39]. Our previous assessments of *C. elegans* food choice behaviors when given the option of six different bacterial diets (OP50/*E. coli* B, HT115/E*. coli K12*, HB101/*E. coli* B-K12 hybrid, *Methylobacterium, Xanthomonas,* and *Sphingomonas*), demonstrates how wildtype (WT) animals are more likely found on any diet except *Methylobacterium.* To determine whether the disinterest in *Methylobacterium* was due to aversion, we conducted pairwise comparisons between *Methylobacterium* and the other five bacterial diets (**Figure 1A**). We measured the choice index of synchronized WT animals whose ancestors (>five generations) were raised on the standard OP50 diet, starting 3-hours after placement on the pairwise assay plate and subsequently at 6-, 24-, and 48-hours. We found that when given a single alternative to *Methylobacterium,* worms were found more often on the other bacterial diet (**Figure 1B-F**). By day 1 of adulthood, each of the pairwise choice indices were positive, indicating that more than half of the animals were found dwelling on any bacterial food source other than *Methylobacterium.* Because parental diet can influence progeny life history traits [2], we repeated this assays but utilized animals derived from parents raised on each diet for more than five generations and found animals still displayed a potent disinterest in *Methylobacterium* (Figures S1A-E).

**Figure 1.**
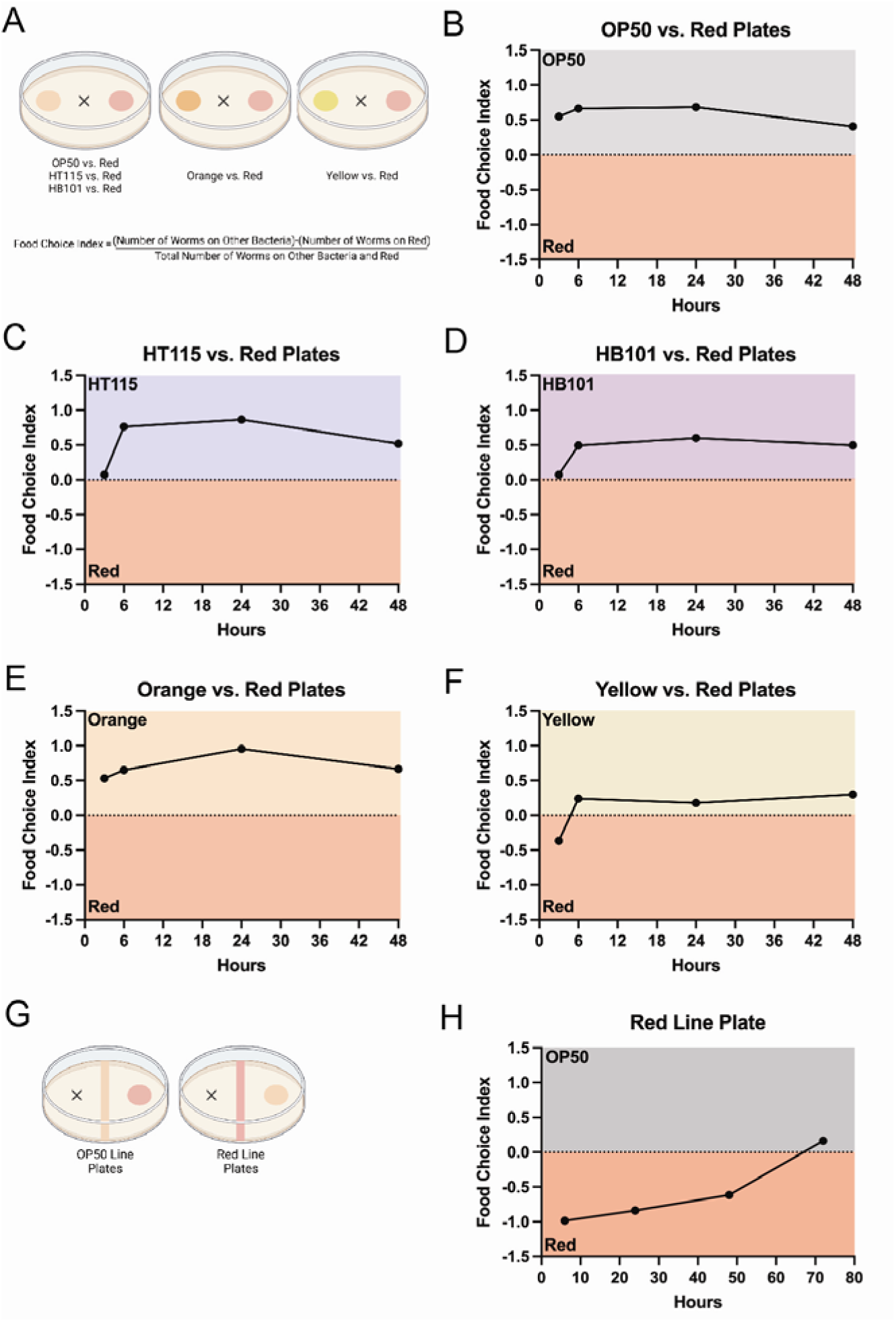
Antipathy behavior of *C. elegans* toward the longevity-promoting *Methylobacterium* diet. (**A**) Schematic for food choice assays and formula used to calculate food choice index. Pairwise food choice comparisons between *Methylobacterium*/Red and OP50 (**B**), HT115 (**C**), HB101 (**D**), *Xanthomonas/*Orange (**E**), and *Sphingomonas*/Yellow (**F**); “x” indicates where animals are placed at start of the assay (0 hours). Experiments were conducted in biological triplicate with three technical replicates. (**G**) Development of forced choice where worms are dropped on the “x” and must interact with one food source before having the option of a difference food source. (**H**) *C. elegans* show antipathy towards *Methylobacterium* and consistently move away from this food source towards *E. coli* until more than half of the population is found on *E. coli* 72 hours after dropping synchronized L1s. Experiments were conducted in biological triplicate with three technical replicates.

A caveat to the two-choice assay design is that the initial choice (the diet that an animal associates with first) limits the capacity to observe changes between the diets over the time course of the experiment when the subject is satisfied with that initial food option. To limit the variation associated with the initial choice, we developed a force choice assay design (**Figure 1G**), which unlike the 2-choice assay design, requires an animal to associate with *Methylobacterium* (red line plate) or *E. coli* (*E. coli* line plate) before having the option to choose the second diet. Similar to the results from the pairwise comparison in the traditional 2-choice assay, wildtype animals dropped on the *E. coli* line plate are found most often on the *E. coli* food; although tracks are found on the *Methylobacterium* spot, indicating animals did explore this diet but returned to the *E. coli* diet (Figure S1F). Intriguingly, when subjected to red line plate assay, after the forced encounter with *Methylobacterium*, wildtype animals are found associated with the second bacterial food source as early as 24 hours (**Figure 1H**) and by 72 hours, more than half of the worms are found off of the red line. Lastly, we tested whether the dislike of *Methylobacterium* in the red line plate was dependent of diet the parental worms were raised on and saw that no matter what food source previous generations of worms are grown on, all display aversion from *Methylobacterium*; as indicated by movement away from *Methylobacterium* and towards *E. coli* in the red line plate assay (Figure S1G). Taken together, these results demonstrate that despite the health and lifespan promoting benefits of the *Methylobacterium* diet [19], wildtype animals consistently prefer any other bacterial food source. Because ancestral exposure to the diet did not influence the antipathy toward *Methylobacterium* in the current generation for either food choice assay, all future experiments were performed with animals derived from OP50-raised parents.

### Sensory neurons influence antipathy behavior away from *Methylobacterium*

To determine the genetic basis of the antipathy response to *Methylobacterium*, we first tested a panel of mutants with defects in neuronal-dependent behaviors[40] (Figure S2A). *dyf-2*, encodes part of the intracellular cilium particle B, that when defective results in a global loss of the ability to detect environmental stimuli[2, 41] and these animals fail to leave the *Methylobacterium* food which revealed the aversion for *Methylobacterium* is a chemosensory response (**Figure 2A**). Surprisingly, a loss-of-function (lf) mutation in *egl-3,* which encodes a proprotein convertase subtilisin/kexin type 2 protein, and a gain-of-function mutation in *egl-30*, a G protein-coupled receptor binding protein, resulted in accelerated movement away from *Methylobacterium* as compared to wildtype animals, indicating that the antipathy leaving behavior for *Methylobacterium* can be modulated (**Figure 2B**). This enhanced distaste for *Methylobacterium* is observed across all three food choice assays paradigms, and the difference can be seen as early as the 24-hour timepoint (Figure S2B-C). Unlike *dyf-2*, which is exclusively expressed in the nervous system, *egl-3* and *egl-30* are expressed in cells outside of neurons and as such, we confirmed that the change in antipathy behaviors were a result of a neuronal function by rescuing wildtype *egl-3* specifically in the nervous system in the *egl-3lf* mutants or driving the expression of the dominant *egl-30gf* allele specifically in neurons of wildtype animals, which reversed or recapitulated the observed antipathy behaviors in response to *Methylobacterium*, respectively (Figure S2D-E). Taken together, these data support the conclusion that antipathy behavior for *Methylobacterium* is controlled by the *C. elegans* nervous system.

**Figure 2.**
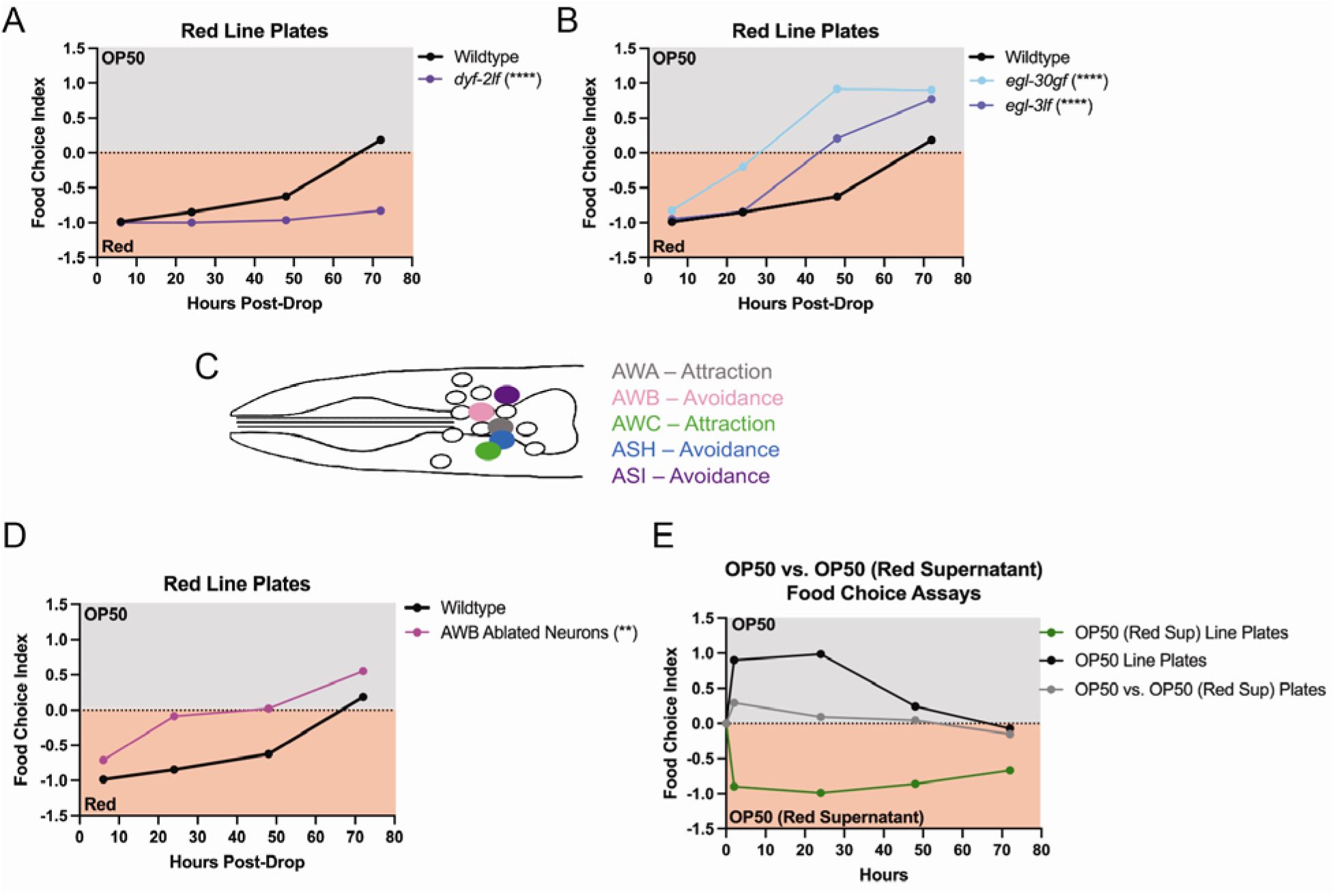
Leaving behavior from *Methylobacterium* is dependent upon chemosensation. (**A**) The inability to detect environmental stimuli in *dyf-2* mutants results in loss of antipathy toward *Methylobacterium* which is in contrast to (**B**) *egl-30gf* and *egl-3lf* mutants that display accelerated movement away from *Methylobacterium*. Significance was calculated at the 72-hour timepoint, p<0.0001 (****). (**C**) Map of the AWA, AWB, AWC, ASH, and ASI chemosensory neuron pairs in *C. elegans* (side view, total of two neurons per pair). (**D**) Genetic ablation of AWB neurons accelerates avoidance from *Methylobacterium,* p<0.01 (**). (**E**) WT animals do not display antipathy for *E. coli* resuspended in *Methylobacterium* growth media.

*C. elegans* chemosensory neurons can detect volatile (olfactory) and water-soluble (gustatory) cues associated with food and other environmental conditions [41]. The amphid sensory organ, which includes eleven pairs of chemosensory neurons each of which responds to specific combinations of molecules, both attractive and repellent [2]. We next tested for changes in preference toward *Methylobacterium* in a panel of chemosensory-defective mutants where individual chemosensory neuron pairs (AWA, AWB, AWC, ASH, ASI) were functionally disrupted (**Figure 2C**). AWB-ablated animals display the strongest antipathy for *Methylobacterium* in the red line plate assay, resulting in almost half the animals observed on the *E. coli* food source at 24 hours – having already moved away from the *Methylobacterium* line first encountered (**Figure 2D** and Figure S2F-H). Similarly, ASH-ablation resulted in ∼50% of animals away from *Methylobacterium* by 48 hours. Loss of *odr-7* results in the inability of AWA neurons to respond to chemoattractant molecules normally sensed by this neuron pair [25], and although not a genetic ablation, these animals have the strongest aversion from *Methylobacterium* by 72-hours with nearly the entire cohort of animals found on *E. coli* (and not on *Methylobacterium*). In addition, *odr-7* mutants remain on *E. coli* more significantly in the 2-choice and *E. coli* line assays. Intriguingly, animals with genetic ablation of ASI neurons were most similar to wildtype in their aversion for *Methylobacterium.* Importantly, we observed that animals did not move slowly; in fact, in several cases animals moved faster as compared to WT worms (Figure S2I-K). As such, the antipathy behavior to *Methylobacterium* is unlikely due to a neuromuscular defect, which the *C. elegans* chemosensory nervous system can influence [2]. Our data reveal a unique type of avoidance response because ASH is critical for the nociception [42], AWB responds to repulsive odorants[ 43–45], and AWA responds to attractant molecules, but ablation of any of these neuron pairs results in enhanced avoidance of *Methylobacterium*, albeit on different time scales. Finally, unlike animals that avoid *Methylobacterium* if given another food choice, WT *C. elegans* did not avoid *E. coli* resuspended in media used to grow the *Methylobacterium* cultures, which suggests that the aversive signal is intracellular and not secreted by *Methylobacterium* into the environment (**Figure 2E**). These findings align with our data showing that the number of animals found on the food in both the *E. coli* line and red line assays at 30 minutes are indistinguishable (Figure S2L); thus, the aversion to *Methylobacterium* occurs after initial contact with the food source. Taken together, this dietary aversion is likely a response to a combination of factors within the *Methylobacterium* diet.

### A *Methylobacterium* diet disrupts *C. elegans* lipid homeostasis

An evolutionarily conserved driver of food aversion is a result of how the diet makes the host feel after eating [46]. One of the most striking phenotypes of animals forced to eat the *Methylobacterium* diet is reduced storage of intracellular lipids[19]. To identify what specific lipid species were most affected we next performed quantitative lipidomics to examine the lipid profiles of WT animals fed the *Methylobacterium* diet as compared to animals fed the standard *E. coli* food source (**Figure 3A-C**, Figure S3A-B, and Table S1). Compared to animals fed *E. coli*, *Methylobacterium*-fed animals had a 3-fold reduction in the TAG/PL ration (**Figure 3A**). We found that the triacylglycerol (TAG) fraction of saturated fatty acids in *Methylobacterium*-fed animals contained higher levels of stearic acid (18:0) and arachidic acid (20:0) (**Figure 3B**). An assessment of the profile of unsaturated fatty acids revealed a 3.4-fold increase in vaccenic acid (18:1) and 2.7-fold more gamma-linoleic acid (18:2) and conversely a significant reduction of several lipid species, including, a 2.9-fold decrease in dihomo-γ-linolenic acid (20:3), a 6.6-fold decrease in alpha-linoleic acid (18:2), and 8.1-fold decrease in oleic acid (18:1) (**Figure 3C**).

**Figure 3.**
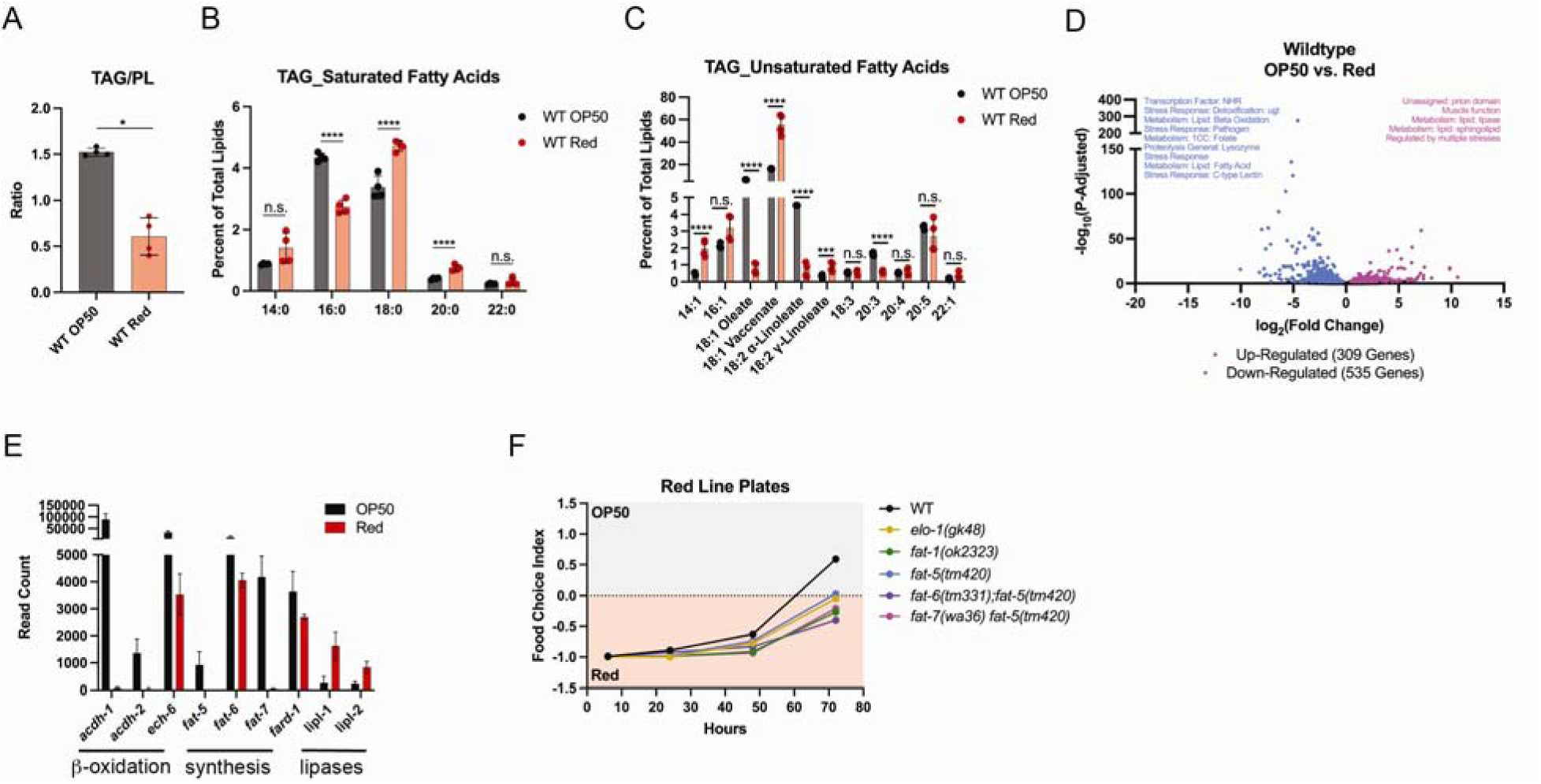
*Methylobacterium* diet alters the landscape of lipid metabolism and storage. (**A**) Relative to animals fed *E. coli*, WT animals fed *Methylobacterium* carry a reduced triglyceride (TAG) to phospholipid (PL) ratio and altered abundance of (**B**) specific saturated lipids and (**C**) unsaturated lipids. (**D**) Transcription signature of WT animals fed *Methylobacterium* reveals display altered expression of lipid homeostasis gene targets (**E**), with the strongest decrease measured in ß-oxidation and lipid biosynthesis genes, but increased expression of lipases. (**F**) Loss of lipid biosynthesis genes can alter antipathy toward *Methylobacterium.* All pairwise comparisons made with Dunnett’s multiple t-test: * p<0.05, ** p<0.01, *** p<0.001, **** p<0.0001.

We next examined changes in the transcriptome of WT animals fed the *Methylobacterium* diet, as compared to an *E. coli* diet by RNAseq, which revealed an enrichment in lipid metabolism regulatory genes (**Figure 3D**, Figure S3C-E and Table S2). We looked carefully at the expression of key lipid homeostasis genes involved in *de novo* lipid biosynthesis (*Fat* and *Elo* family;), lipid transport (Vitellogenin family), lipid storage (e.g., *dgat-2*, *seip-1*, *dhs-3*, *acs-4*, *atgl- 1*), and lipid degradation (fatty acid β-oxidation, *Acdh* and *Ech* family), which revealed several significant changes (**Figure 3E** and Table S2). Although a modest (∼1.5-fold) increase in *fasn-1* was measured, in general, the expression of all *Fat* and *Elo* genes was markedly reduced with a striking 85-fold and 167-fold reduction in the expression of *fat-7* and *fat-5*, respectively, being the most significant changes detected.

Because endogenous lipid profiles are linked to the activity of specific lipid biosynthesis enzymes [47], we tested if the changes in the expression of lipid biosynthesis genes was linked to the antipathy toward the *Methylobacterium* diet. Informed by the lipidomic and transcriptional analyses, we tested animals with mutations in the lipid biosynthesis enzymes that metabolize the species differentially impacted by the *Methylobacterium* diet (**Figure 3F** and Figure S3F). At 72 hours, >50% of WT animals have moved away from the *Methylobacterium* and are found on the *E. coli* diet, but strikingly *elo-1lf*, *fat-1lf*, and *fat-5lf* single mutants and double mutants of *fat- 6lf*;*fat-5lf* and *fat-*7*lf*;*fat-5lf* display a diminished antipathy toward *Methylobacterium*. Collectively, these data reveal that disruption of the homeostatic balance of endogenous lipid species can influence the initial response to the *Methylobacterium* diet, but also alter the response to the diet after prolonged exposure.

### Diets with high saturated fat evoke antipathy behaviors through vaccenic acid

The macro- or micronutrient composition of bacteria can be vastly different [42, 48] and we hypothesized that the aversion to *Methylobacterium* could be a response to dietary metabolites. We performed untargeted mass spectrometry of the six bacterial diets to discover metabolites with differential abundance and molecules unique to *Methylobacterium* (**Figure 4A** and Figure S4A-B, and Table S3). With the exception of the OP50 and HB101 *E. coli* food sources which display similar the metabolic profiles, each of the other diets were distinct (Figure S4A-D). Relative to the standard OP50 *E. coli* diet, we found eight metabolites that were significantly different in *Methylobacterium* as compared to the other five bacterial food sources (Figure S4E, and Table S3). Three of these *Methylobacterium* enriched metabolites, salicylic acid, valeric acid, and palmitic acid were greater than 5-fold higher in *Methylobacterium* as compared to *E. coli*. In a tradition 2-choice assay, over the 72-hour time course, animals are free to move between the two diet choices. We next supplemented *E. coli* with a 40mM dose of each metabolite uniquely enriched in *Methylobacterium* and discovered that wild type animals displayed a strongly aversive response to the presence of excess palmitic acid - nearly all animals avoided the supplement - but not valeric acid or salicylic acid which consistently had ∼50% of the population on each food spot (**Figure 4B**). These data suggest that *C. elegans* can sense the presence of excess lipids. Palmitate is the building block for nearly all lipid species in *C. elegans* acting as a precursor substrate and building block for the diverse and more complex lipids that can be synthesized *de novo* (**Figure 4C**). We next tested whether the lipid products generated by the lipid biosynthesis enzymes that influenced antipathy toward *Methylobacterium* could evoke leaving behaviors in a modified forced choice assay where WT animals first encounter OP50 supplemented with vehicle (EtOH) or a specific fatty acid. Recalling that the loss of *fat-5* attenuated the leaving behavior from *Methylobacterium* (**Figure 3** and Figure S3), we first tested supplementation with vaccenic acid (**Figure 4D**), which is the elongated product of palmitoleate (16:1) generated by the FAT-5 delta-9 desaturase, that evoked a strong leaving response. This response was specific as supplementation with oleate, the direct product of FAT-6 and FAT-7 delta-9 desaturases and downstream product of ELO-1 elongation of palmitate, evoked an avoidance response comparable to vehicle supplementation (Figure S4F). These data reveal that lipids in the diet and specific intermediates of endogenous lipid metabolism are key signals in the decision matrix to avoid a particular diet recently ingested.

**Figure 4.**
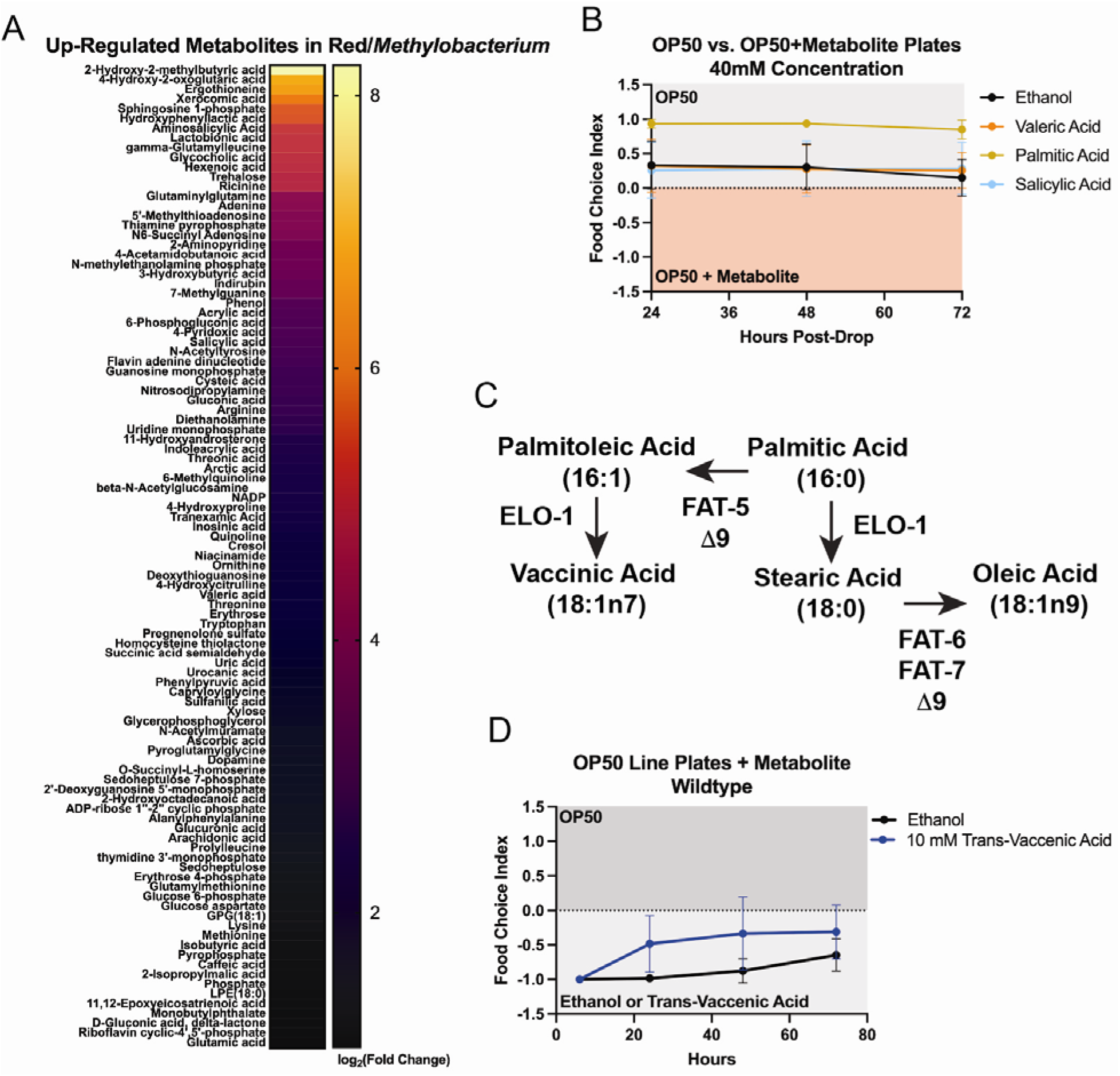
Lipids enriched in the *Methylobacterium* diet influence antipathy behavior. (**A**) Metabolite profiling of *Methylobacterium* identifies several fatty acids (FA) that are higher in content in *Methylobacterium* in comparison to *E. coli/*OP50. (**B**) *C. elegans* display enhanced avoidance of *E. coli* supplemented with palmitic acid (16:0 FA), but not valeric acid (5:0 FA) or salicylic acid (cyclic organic). (**C**) Vaccenic acid, which can be made from palmitic acid via the enzymatic activities of FAT-5 and ELO-1, drives antipathy-like behavior in WT animals when supplemented to the *E. coli* diet (**D**).

### The sphingosine rheostat regulates the response to *Methylobacterium*

To better define the signaling events between ingestion of the *Methylobacterium* diet and AWB/AWC signaling of aversion, we returned to our metabolomic and transcriptomic data (**Figure 3**, Table S1-2). Although saturated fats were uniquely in excess in the *Methylobacterium* diet, several additional metabolites linked to sphingosine metabolism were found in *Methylobacterium* and at least one other diet, as compared to *E. coli*. The sphingosine rheostat balances the abundance of Sphingosine-1-phosphate (S1P) which is a bioactive lipid mediator, derived from ceramide (**Figure 5A**). We found S1P levels were 5.5-fold more abundant in the *Methylobacterium* diet as compared to *E. coli* and correlatively, sphingosine, the precursor of S1P, was reduced, relative to *E. coli* (Figure S5A and Table S3).

**Figure 5.**
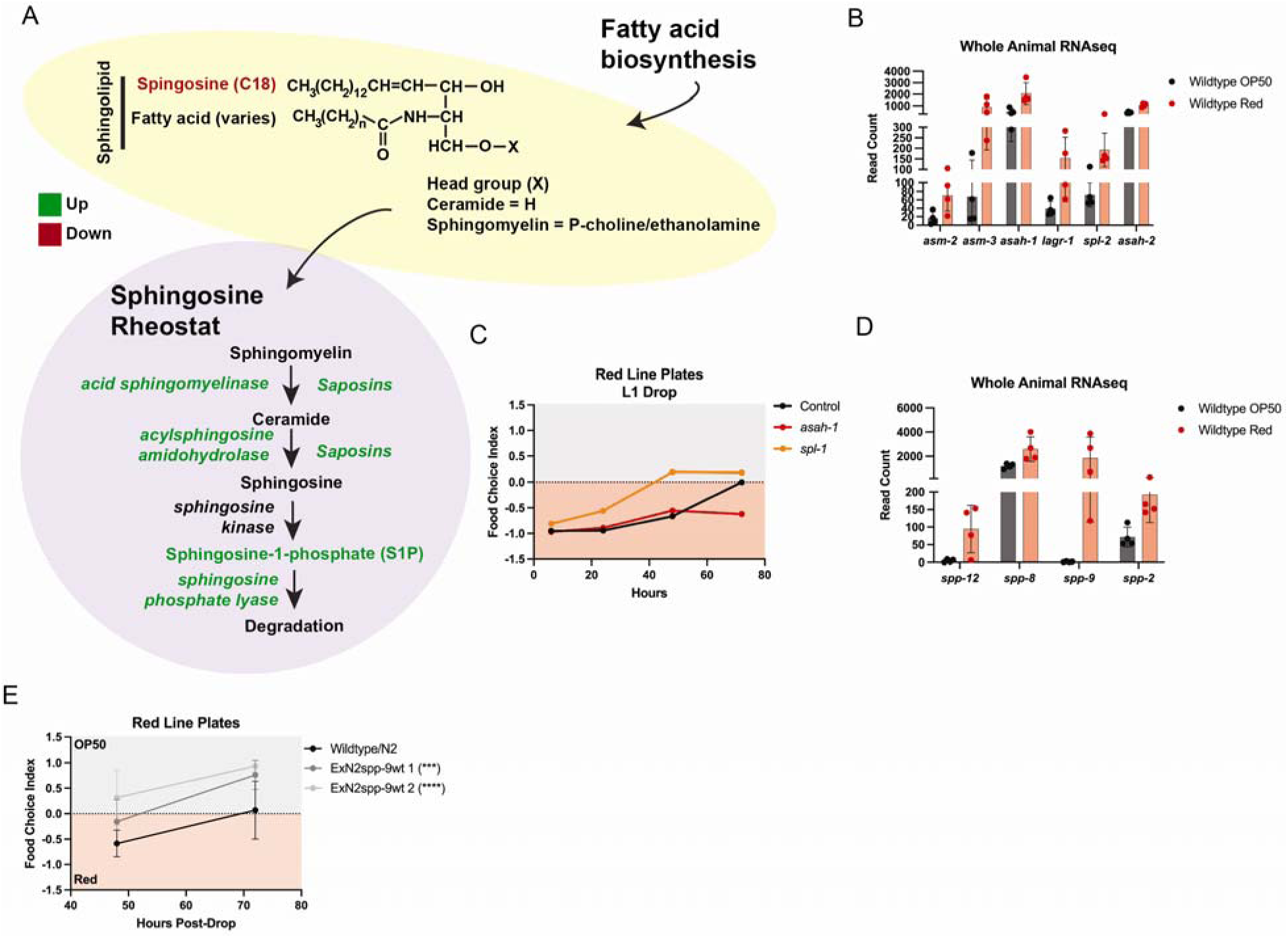
Sphingosine-1-phosphate (S1P) mediates antipathy behavior toward *Methylobacterium.* (**A**) Cartoon depiction of the sphingosine rheostat. (**B**) WT animals fed *Methylobacterium* display increased expression of genes encoding enzymes that function in the sphingosine rheostat. (**C**) Antipathy behavior can be enhanced or impaired by altering the expression of rheostat genes *spl-1* and *asah-1* by RNAi, respectively. (**D**) The expression of several saposin-related genes, that generate sphingosine from ceramide and sphingomyelin, display increased expression in WT animals fed *Methylobacterium.* (**E**) Overexpression of *spp- 9* drives enhances antipathy toward *Methylobacterium*, resulting in significance from WT at 72 hours post-drop. All pairwise comparisons made with Dunnett’s multiple t-test: * p<0.05, ** p<0.01, *** p<0.001, **** p<0.0001.

Among the genes most responsive to the *Methylobacterium* diet were genes involved in the sphingosine rheostat, including: a 2.27 and 6.07 log_2_-fold increase in the expression of the acid sphingomyelinase genes *asm-2* and *asm-3*, respectively; a 1.81 and 1.25 log_2_-fold increase in the acylsphingosine amidohydrolase genes *asah-1* and *asah-2*, respectively, which generates sphingosine from ceramide; and a 1.32 log_2_-fold increase in the sphingosine phosphate lyase gene *spl-2*, respectively, which can irreversible degrade S1P (**Figure 5B** and Figure S5B).

Although sphingosine metabolism is essential for normal development in *C. elegans* [49, 50], to determine causality for the changes in sphingosine signaling, we tested the importance of these transcriptional changes to the antipathy behaviors of WT worms toward the *Methylobacterium* diet by RNAi. Strikingly, RNAi reduction of *spl-1* expression accelerated leaving behavior from *Methylobacterium* while reducing *asah-1* expression delayed leaving (**Figure 5C** and Figure S5C). Because loss of *asah-1* would reduce the generation of S1P by impeding ceramide synthesis while loss of *spl-1* would increase S1P levels, these data suggest that S1P signaling promotes antipathy behaviors of WT animals toward *Methylobacterium*.

Another interesting class of genes related to the sphingosine rheostat that display differential expression on the *Methylobacterium* diet are the Saposin-like type B family of proteins, specifically *spp-12, spp-9, and spp-2* (**Figure 5D**). Saposins and saposin-like proteins are evolutionarily conserved small lipid interacting proteins, that are essential for sphingolipid degradation and membrane digestion, which can generate sphingosine [51, 52]. In light of the significant increase in the expression of *spp-9* in WT animals fed the *Methylobacterium* diet, we tested the overexpression of *spp-9* in WT animals which in support of our model where S1P drives antipathy and leaving of the *Methylobacterium* diet, overexpression of *spp-9* induced enhanced/accelerated leaving from *Methylobacterium* (**Figure 5E**). Taken together, these data support S1P signaling in the antipathy for *Methylobacterium* which can be modulated to alter the strength of this behavioral response.

### *odr-1* signaling is essential for the antipathy behavior toward *Methylobacterium*

Based on the observed involvement of the sensory nervous system for the antipathy behavior of WT animals exposed to *Methylobacterium*, we next tested a panel of genetic mutants with specific defects in chemosensation for altered aversion for *Methylobacterium* and identified mutants with enhanced (early leaving within 24-hours) and four mutants that failed to display antipathy toward *Methylobacterium.* Among the genetic mutations that fail to leave *Methylobacterium* were the guanylate cyclase *odr-1lf*, *daf-11lf*, *che-2lf*, and *lim-4lf* (**Figure 6A** and Figure S6A-C); however, *lim-4lf* animals have movement defects which likely confound the leaving behavior (Figure S6D). *odr-1* is expressed in AWB and AWC neurons, but not AWA [53, 54]. Importantly, the reduced leaving behavior from the *Methylobacterium* food was specific to loss of functional *odr-1* in these neuron pairs as ectopic expression of WT *odr-1* specifically in the AWB neuron pair under the *str-1* promoter (**Figure 6B**) or in a single AWC neuron under the *str-2* promoter [44, 55] in the *odr-1lf* mutant not only restored the antipathy leaving behavior resulting from loss of *odr-1*, but significantly enhanced leaving as compared to the normal antipathy response in WT animals (**Figure 6C**).

**Figure 6.**
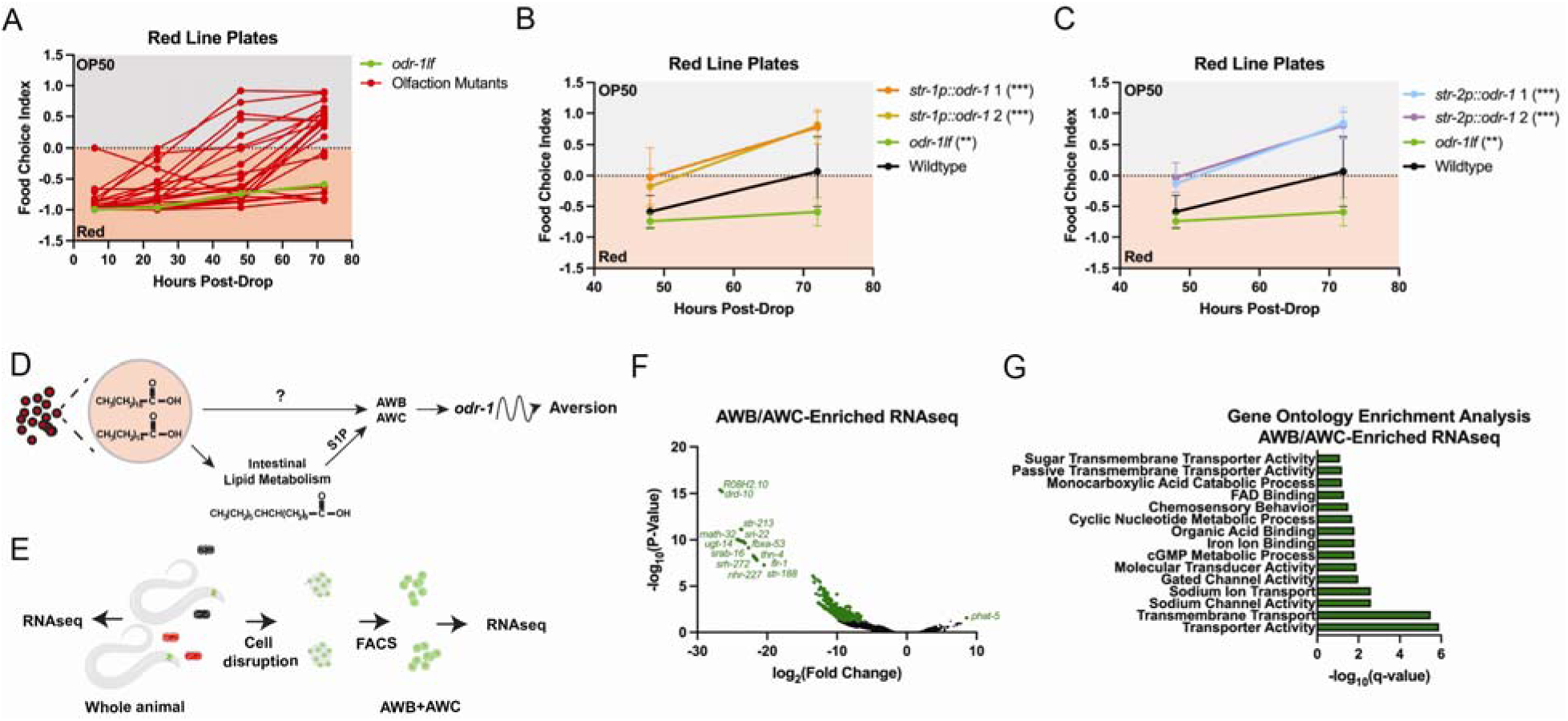
ODR-1 signaling is required for the antipathy response towards *Methylobacterium.* (**A**) *odr-1lf* mutants don’t move away from *Methylobacterium* at any point from L1-Day 1 of adulthood on Red Line Plates. Rescuing *odr-1* in AWB neurons only (**B**) or AWC neurons only (**C**) in the *odr-1lf* mutant restores antipathy behavior towards *Methylobacterium*. (**D**) Cartoon depiction of how *C. elegans* sense high saturated fats provided by *Methylobacterium*, leading to the avoidance phenotype due to signaling through AWB and AWC neurons. (**E**) Schematic for AWB and AWC neuronal isolation using *odr-1p*::RFP for bulk AWB/AWC-enriched RNAseq. (**F**) AWB/AWC-enriched RNAseq revealed all but one gene (*phat- 5)* is down-regulated in neuron populations in response to *Methylobacterium*. (**G**) Gene ontology enrichment identified transporter activity, transmembrane transport, and chemosensory behaviors as top changes in response to exposure to *Methylobacterium*. All pairwise comparisons made at 72 hours post-drop with Dunnett’s multiple t-test: * p<0.05, ** p<0.01, *** p<0.001, **** p<0.0001.

We next conducted an RNAseq analysis of the *C. elegans* transcriptional response to *Methylobacterium* across organism and cellular hierarchies. We first examined the differential transcriptional response to the *Methylobacterium* diet in *odr-1lf* mutants as compared to WT animals [19]. GO-term/Enrichment/ and analyses of the context-specific (diet) changes in transcripts between *odr-1lf* mutants revealed significant changes in stress responses and lipid homeostasis, similar to that observed in WT animals (Figure S6E), in fact very few differences were found in the transcriptional response to the *Methylobacterium* diet between WT and animals lacking functional *odr-1*(Figure S6F); including the enhanced expression of genes involved in sphingosine metabolism(Figure S6G-I). These data suggest the role *odr-1* plays in AWB/AWC is downstream of the induced *spp-9* transcripts, which normally are most highly expressed in the intestine[56] (**Figure 6D**), which fits our model suggesting the antipathy response occurs after the *Methylobacterium* diet is ingested.

Finally, because RNA isolation from whole animals can make detection of transcriptional responses from specific cells difficult to measure, we next utilized a strain harboring an integrated *odr-1p::RFP* transgene, which only labels the AWB and AWC neuron pairs [57], to enrich AWB and AWC neurons by FACS from animals fed *E. coli* or *Methylobacterium* to compare the specific transcriptional response in *odr-1* expressing sensory neurons (**Figure 6E** and Table S4). Gene enrichment analysis revealed differential expression of genes with functional roles in cellular signaling, metabolism, and chemosensory behaviors (**Figure 6F-G** and Table S4). These data reveal the AWB and AWC neurons are sensitive and transcriptionally responsive to the *Methylobacterium* diet.

### Antipathy behavior impedes the longevity-promoting effects of the *Methylobacterium* **diet**

To test the association, if any, between the antipathy behavior that WT animals have for the *Methylobacterium* diet and the longevity response observed when it is provided as the only diet option, we measured the lifespan of WT, *odr-1lf*, and animals with AWA, AWB, or AWC ablated neurons on the longevity-promoting *Methylobacterium* diet as compared to the standard *E. coli* OP50 diet. The *Methylobacterium* diet was capable of extending the lifespan in all genetic backgrounds (**Figure 7A-E** and Table S5). Whereas the *Methylobacterium*-induced increase in lifespan was maintained in *odr-1* mutant (**Figure 7B**) and AWA signaling-deficient animals (**Figure 7C**), remarkably, the hazard ratio for animals eating *E. coli* versus *Methylobacterium* was 2.1 times greater for animals lacking AWB signaling (**Figure 7D**) and 1.3 times greater for animals without functional AWC signaling (**Figure 7E**) as compared to WT animals. These data functionally uncouple the longevity-effects of the *Methylobacterium* diet from the antipathy behaviors and more importantly reveal that in the absence of leaving behavior (AWB and AWC ablation), animals will spend more time on the *Methylobacterium* diet and benefit from a further increase in lifespan. These results suggests that the AWB and AWC neuroendocrine circuit mediates behaviors that cause avoidance of a diet that is generally beneficial for their overall health across the lifespan (**Figure 7F**).

**Figure 7.**
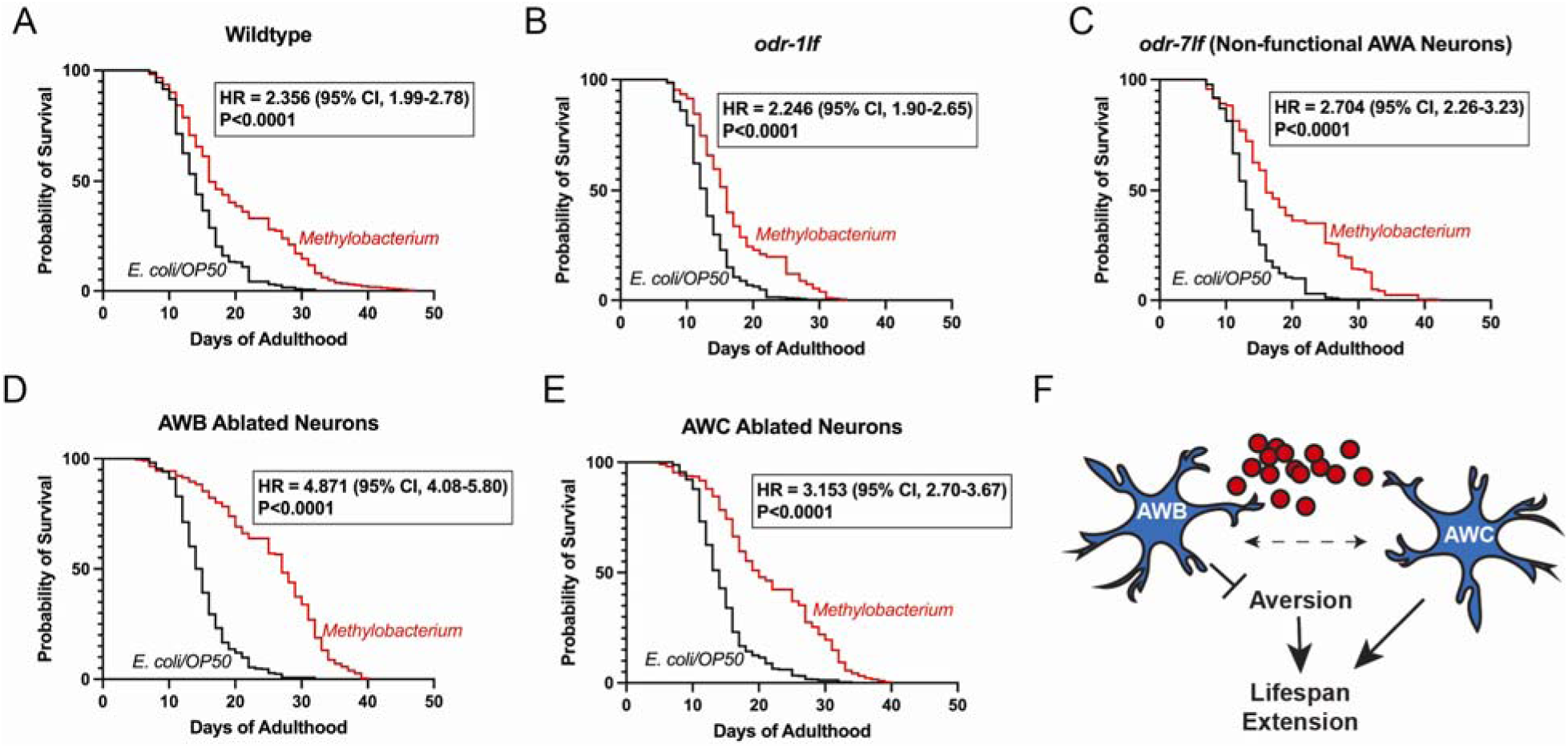
AWB and AWC neurons impede the longevity response from the *Methylobacterium* diet. (**A**) Wildtype (**B**) *odr-1lf* mutants, (**C**) mutants with non-functional AWA neurons, (**D**) AWB-ablated neuron mutants, and (**E**) AWC-ablated neuron mutants all exhibit diet-dependent lifespan extension in the presence of *Methylobacterium*. P-value was determined using log-rank (Mantel-Cox) test for survival. (**F**) Cartoon schematic highlighting the importance of AWB and AWC sensory neurons in an avoidance response to the lifespan- promoting *Methylobacterium* diet.

Taken together, these data support a behavioral program where animals integrate nutritional information to make dietary choices when more than one option exists. The decision to eat *E. coli* or *Methylobacterium* involves, at least in part, an assessment of endogenous and dietary lipids and signaling through ODR-1 expressing AWB and AWC sensory neurons that mediate an aversive behavioral response. This discovery provides a new framework for understanding how the microbiome [58], as a diet for *C. elegans*, can influence perceptions of diet quality and decisions on what food to eat.

## DISCUSSION

Diet-gene interactions have the potential to influence nearly all facets of health, and diet remains one of the most variable aspects of health between individuals and as such, understanding the complex relationship between genetic variation, diet composition, and diet availability is difficult. In the wild, decisions about what food to eat require a value judgement that weighs nutritional needs, value, past experience, and safety. The cellular outcomes that are influenced by diet with age are malleable and influenced by functional capacity, energetic reserves, and responsiveness to treatment [17, 59, 60]

Previous work has shown that the presence of different bacterial communities in the *C. elegans* natural environment can shape multiple aspects of *C. elegans* biology including changes in physiological attributes over the lifespan including metabolic homeostasis, reproductive capacity, gene expression profiles, and behavior [19].

Chemosensory mechanisms have been studied immensely, and clearly demonstrate that volatile drive to an attractive or aversion response [3]. *C. elegans* not only pick up on these cues but use them to learn how to avoid future interactions with toxic and pathogenic microbes and thus improve their chances for better health and survival [7, 61, 62]. Intriguingly, evidence suggests that *C. elegans* choose bacterial diets that are more beneficial for growth and longevity[2, 39]. Our work challenges this model as *Methylobacterium* promotes precocious development, metabolic refinement, stress resistance and extended lifespan, but wildtype animals choose any bacterial diet over *Methylobacterium* – even at the expense of health.

Dietary behaviors are often measured by giving animals a choice of two or more options, but these methods are confounded by the organisms initial choice of diet. Because the antipathy behavior is a leaving response, we developed new forced choice assays where animals must interact with one bacteria first making the measurement of leaving possible to quantify. These assays revealed that worms actively move away from *Methylobacterium* and towards other bacterial diets and this is not just a random or passive response.

Inspired by previously described food choice studies that demonstrated food choice can be due to chemosensory [41, 63] or gustatory signals that respond to attractive and aversive chemicals [64], we disrupted these pathways with an amalgam of mutants that disrupt this neuronal signaling and saw a spectrum of responses to *Methylobacterium*. Our work supports the idea that the avoidance of *Methylobacterium* is determined, at least in part, by the ciliated amphid neurons of the *C. elegans* brain. Importantly, the antipathy response can be modulated as mutations that regulate activity can influence the intensity of avoidance or even attraction to *Methylobacterium.* Although this finding is critical to understanding the antipathy response, what is perhaps most surprising is that the avoidance behavior to *Methylobacterium* requires ingestion of the microbe at which point the host then leaves the bacteria in search of something better. Host-microbe interactions can also promote changes in cellular function [65] and therefore dose responses to a particular microbial environment might regulated and the level of behavior to control the magnitude of a response.

We expected that the source of antipathy in the *Methylobacterium* diet would be a toxic xenobiotic, but our metabolic profiling and testing of specific molecules enriched in the *Methylobacterium* cells revealed that specific lipids could act as potent drivers of leaving behavior. Our genetic dissection of this response, in concert with dietary supplementation of specific lipid species, revealed that saturated lipids and the metabolic derivatives, vaccenic acid, are sufficient to promote leaving behaviors once ingested. These results suggest that the current state of lipid homeostasis and more importantly the abundance of particular lipids ingested can influence an animals decision-making process for what food to eat and continue eating.

Beyond their utility for energy storage and fuel, lipids are critical molecules for cellular structure, function, and signaling. The sphingosine rheostat centers around sphingosine-1-phosphate (S1P), a bioactive lipid mediator that regulates immunomodulation. RNAseq analysis of animals fed *Methylobacterium* reveals that animals alter the expression of enzymes that influence the sphingosine rheostat, perhaps in an attempt to establish homeostasis. Ultimately, feeding *Methylobacterium* increases S1P and genetic manipulations that increase S1P (loss of sphingosine phosphate lyase or increased saposin expression) enhance the antipathy response. Similarly, interventions that decrease S1P synthesis, such as *asah-1* RNAi, make animals blind to *Methylobacterium*. Taken together, it is evident that S1P signaling is important to the response to *Methylobacterium* which when altered can modulate food-related behaviors and supports its potential use as a pharmacological agent [66].

We tested whether the extended lifespan associated with feeding the *Methylobacterium* diet was influenced by the same neuroendocrine signaling pathways that rule the antipathy response. Ablation of AWB neurons enhances the antipathy response, but when animals without AWB signaling are put on plates where *Methylobacterium* is the only food source, and avoidance isn’t possible, the increase in lifespan is even greater. Similarly, loss of AWC neurons also enhances the longevity response to the *Methylobacterium* diet but does not impact antipathy behavior. AWB and AWC share ODR-1 signaling and *odr-1lf* mutants do not display antipathy for *Methylobacterium* and similarly do not benefit from the additive lifespan response to *Methylobacterium*. These data suggest that AWB and AWC neurons integrate *odr-1* signaling in response to *Methylobacterium,* but more importantly, the ability to distinguish *Methylobacterium* and avoid it is important for longevity.

In an age where individuals can design the composition of their microbiome with the goal of improving health [9, 30, 31, 58], our study reveals genetically coded antipathy behavioral program where after trying a diet, animals integrate nutritional information from that food source to make choices about whether they should continue eating or seek out other options. The process that individual animals use when making the choice between *Methylobacterium* and *E. coli* integrates ODR-1 signaling in AWB and AWC sensory neurons, the sphingosine rheostat, and an assessment of the current lipid storage landscape. Importantly, this work reveals the complex decision matrix utilized to make dietary preference decisions based on contemporaneously available data on the current metabolic state of the organism and the nutritional composition of the food.

## ACKNOWLEDGEMENTS

We thank S. Ledgerwood for technical assistance. This work was funded by the NIH R01AG058610 and Hevolution Foundation award HF AGE-004 to SPC, F31AG077873 to NLS, F31GM137587 to CDT, and T32AG052374 to NLS and CDT. We also thank the USC School of Gerontology Imaging Core that is funded in part by the Nathan Shock Center of Excellence P30AG068345. Some strains were provided by the CGC, which is funded by the NIH Office of Research Infrastructure Programs (P40 OD010440). We thank WormBase for database curation and data access.

---

**Figure S1.**
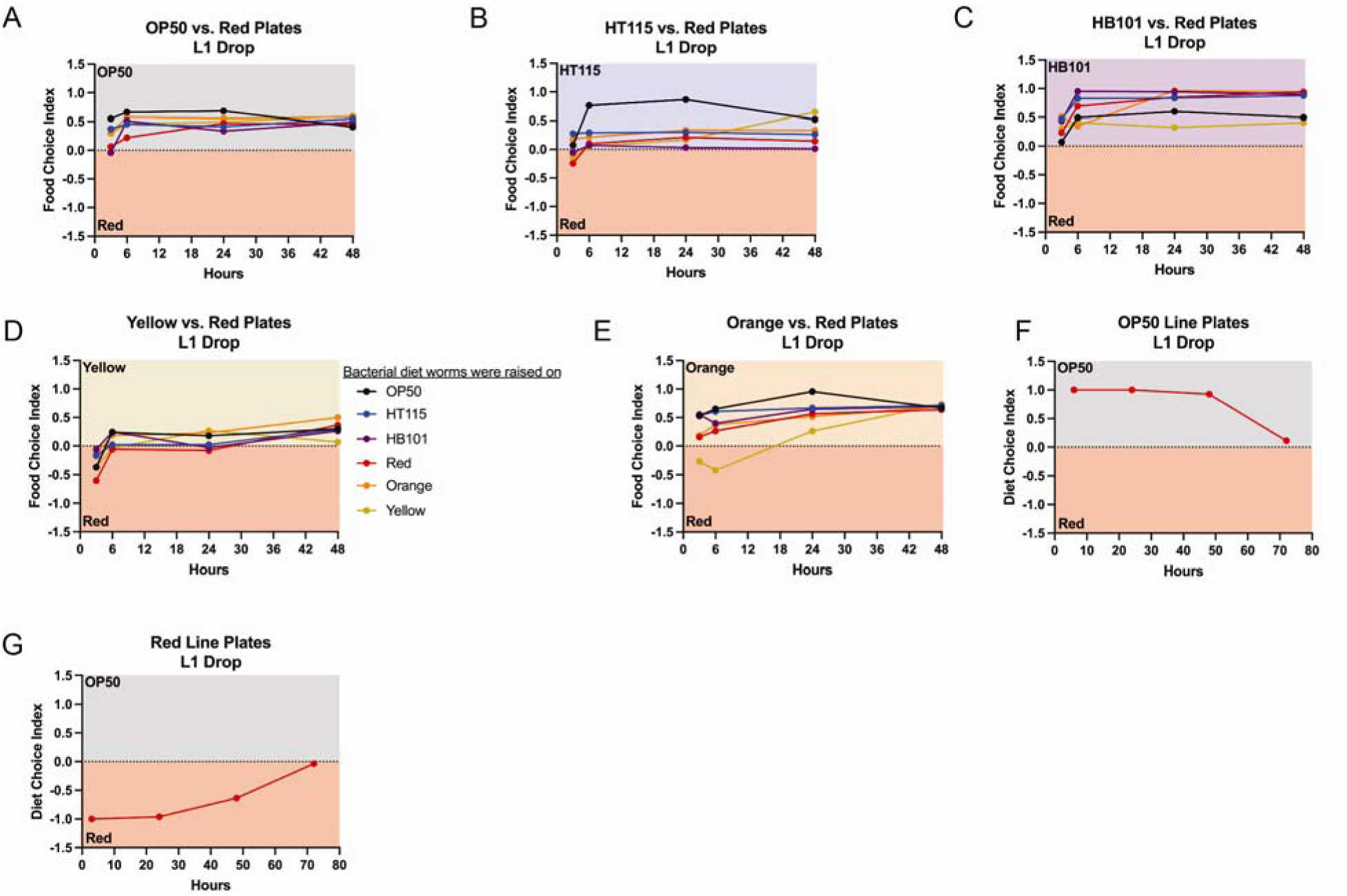
*Methylobacterium* leads to the strongest avoidance response compared to other bacterial diets in a diet-independent manner. When *C. elegans* are given the option of either *Methylobacterium* or another bacterial diet, worms will be found more often on the other bacteria. Worms prefer OP50 (**A**), HT115 (**B**), HB101 (**C**), *Sphingomonas*/Yellow (**D**), and *Xanthomonas/*Orange (**E**) over Red/*Methylobacterium*. More sophisticated assays using an OP50 Line Plate (**F**), and Red Line Plate (**G**) reveal more occupancy on OP50/*E. coli*.

**Figure S2.**
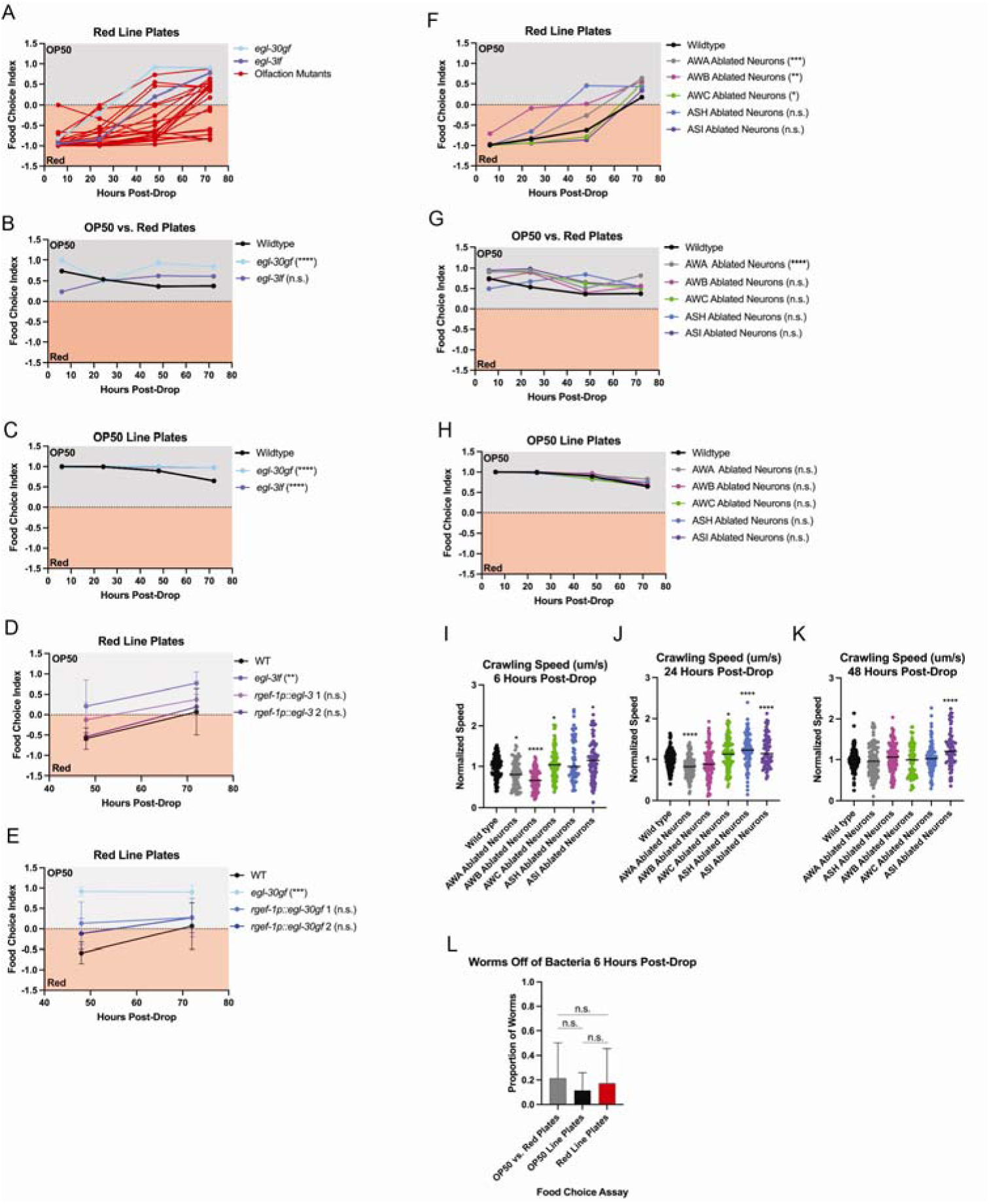
Chemosensation in *C. elegans* mediates the avoidance response to *Methylobacterium*. Of the olfaction mutants tested, (**A**) *egl-3lf* and *egl-30gf* mutants move faster away from the Red/*Methylobacterium*. *egl-30gf* and *egl-3lf* are found more often on OP50 than wildtype worms on both OP50 vs. Red Plates (**B**) and OP50 Line Plates (**C**). On Red Line Plates, rescuing *egl-3lf* pan-neuronally using the *rgef-1p::egl-3* construct returned food choice back to wildtype (**D**). Expressing *egl-30gf* only pan-neuronally using the *rgef-1p::egl-30gf* construct (**E**), also returned food choice back to wildtype levels. (**F**) Ablated chemosensory neuronal mutants displayed different ranges of food choice behavioral changes on Red Line Plates (**F**), OP50 vs. Red Plates (**G**), and OP50 Line Plates (**H**). All pairwise comparisons made at 72 hours post-drop with Dunnett’s multiple t-test: * p<0.05, ** p<0.01, *** p<0.001, **** p<0.0001. (**I-K**) Crawling speed of neuronal ablation mutants at each timepoint of food choice assay. Comparisons made to wildtype worms using Dunnett’s multiple t-test: * p<0.05, ** p<0.01, *** p<0.001, **** p<0.0001. (**L**) Regardless of the food choice assay, more worms are found on food at the 6-hour timepoint post-drop.

**Figure S3.**
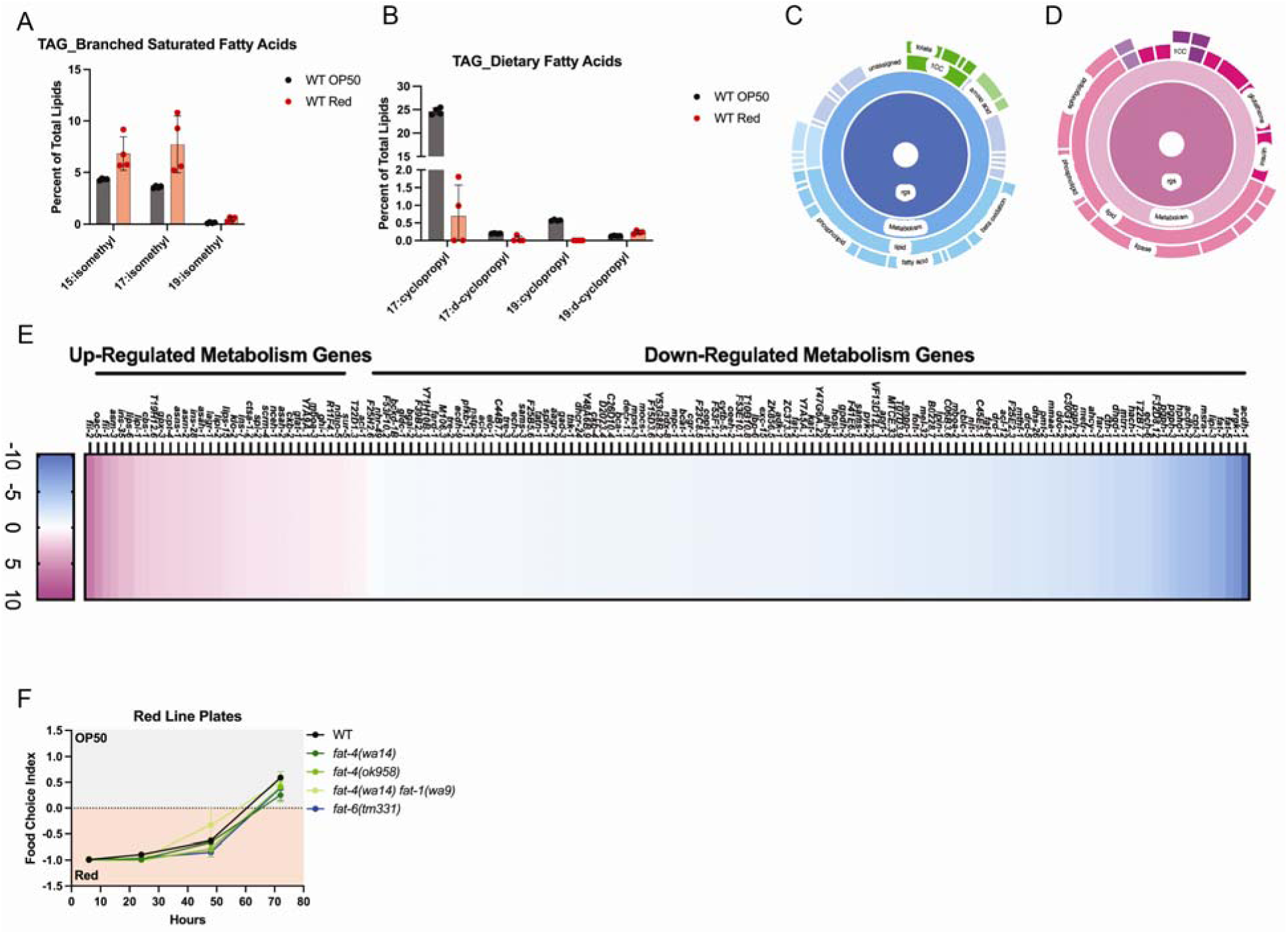
Fatty Acid Metabolism is altered in worms fed OP50/*E. coli* versus *Methylobacterium.* (**A**) Wildtype worms fed Red/*Methylobacterium* have higher levels of 15:isomethyl and 17:isomethyl. (**B**) *Methylobacterium* provides far less dietary fatty acids compared to OP50/*E. coli*. (**C-E**) RNAseq analysis revealed lipid metabolism to be one of the main gene ontologies effected by worms fed *Methylobacterium*. (**F**) Loss-of-function mutations in *fat-4, fat-1,* and *fat-6* do not change food choice preference on Red Line plates.

**Figure S4.**
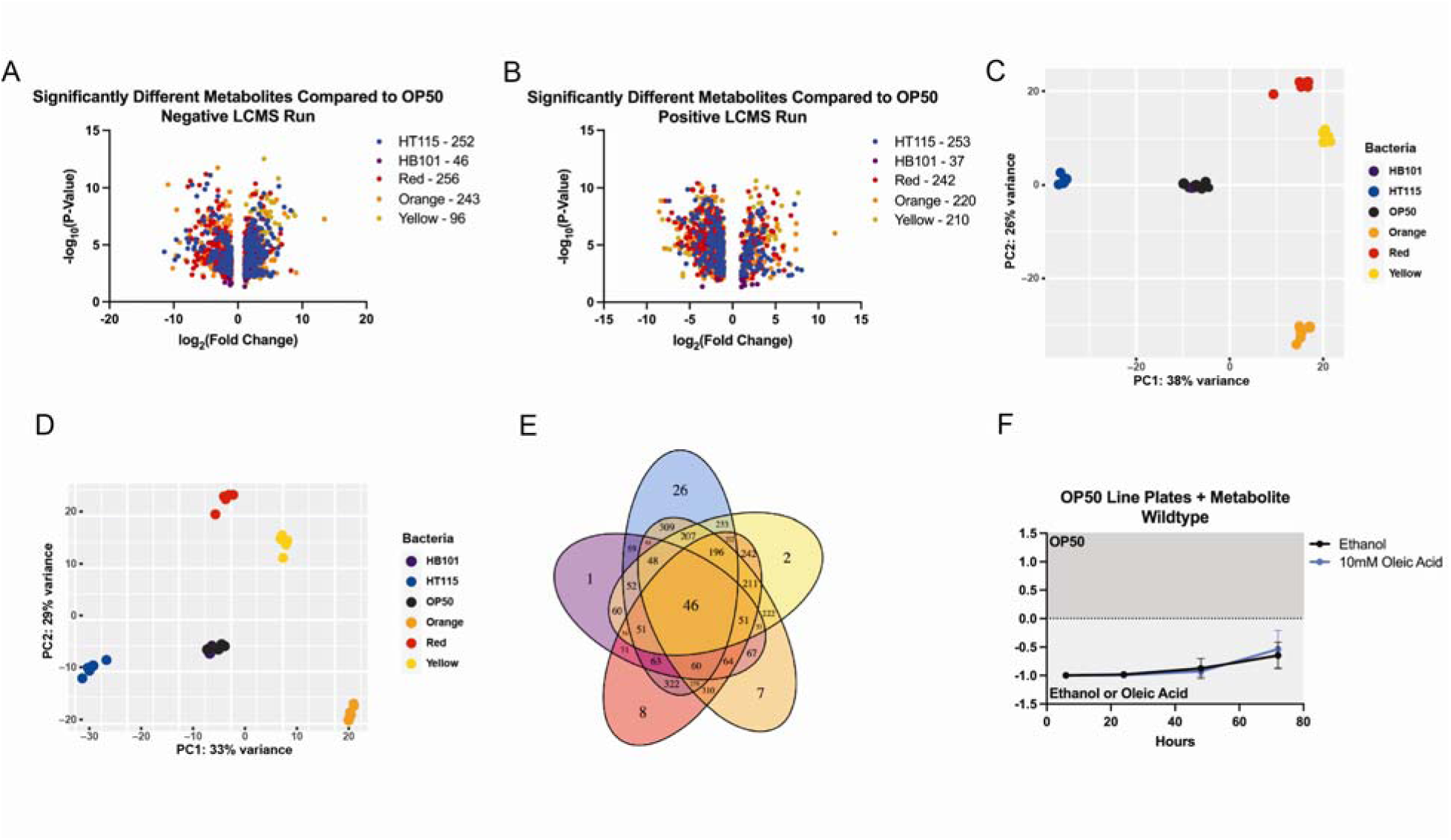
Bacterial diets supply *C. elegans* feeding on them different contents of metabolites. (**A-B**) Significantly different metabolites found in all five bacterial genera compared to *E. coli*/OP50 in both the negative (**A**) and positive (**B**) LCMS runs. The number next to the bacterial diet is the number of differently regulated metabolites in each bacterial diet (p<0.05). (**C-D**) PCA plots reveal each bacterial diet metabolite content is distinctly different from *E. coli*/OP50 except that of *E. coli/*HB0101. (**E**) Venn diagram showing the overlaps in differentially regulated metabolites in each bacterial diet. HT115 (blue) has 26 unique metabolites, HB101 (purple) contains 1 unique metabolite, *Methylobacterium/*Red (Red) contains 8 unique metabolites, *Xanthomonas*/Orange (Orange) contains 7 unique metabolites, and *Sphingomonas*/Yellow (Yellow) contains 2 unique metabolites. (**F**) Wildtype worms do not avoid oleic acid when present on an OP50 Line Plate.

**Figure S5.**
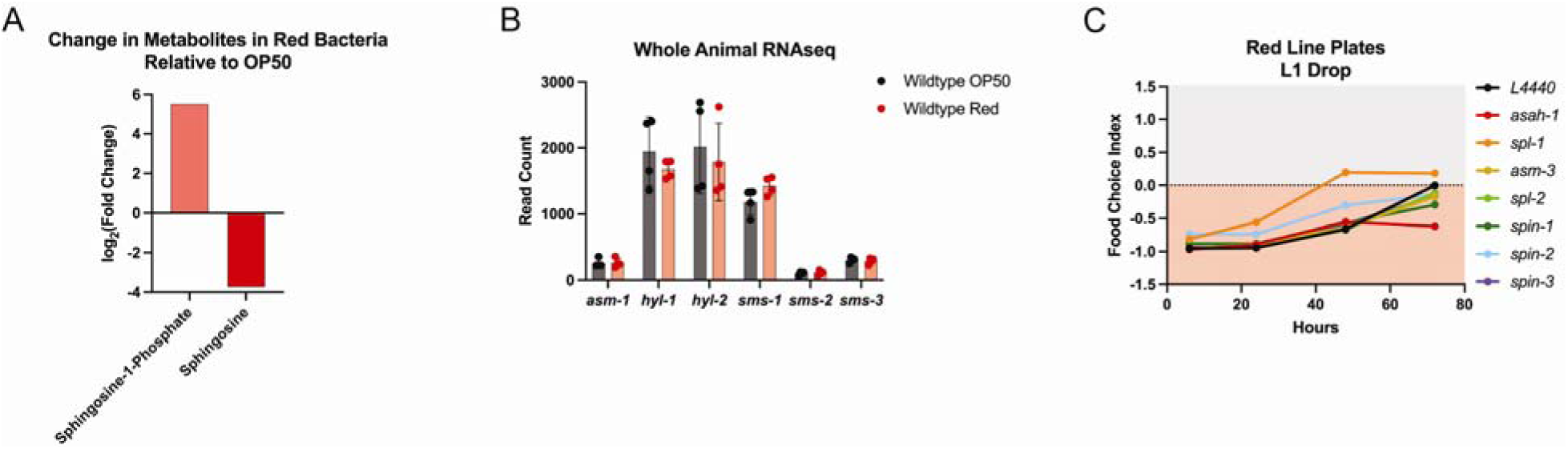
*Methylobacterium* regulates sphingosine metabolism through dietary content and altering of gene expression in sphingosine rheostat pathways. (**A**) *Methylobacterium* contains higher levels of sphingosine-1-Phosphate (S1P), but lower levels of sphingosine compared to *E. coli/*OP50 as measured by untargeted metabolomics. (**B**) Wildtype worms fed *E. coli*/OP50, and *Methylobacterium*/Red contain similar gene expression read counts in sphingosine metabolism pathways. (**C**) RNAi of genes in sphingosine metabolism pathways alters avoidance response of *Methylobacterium*.

**Figure S6.**
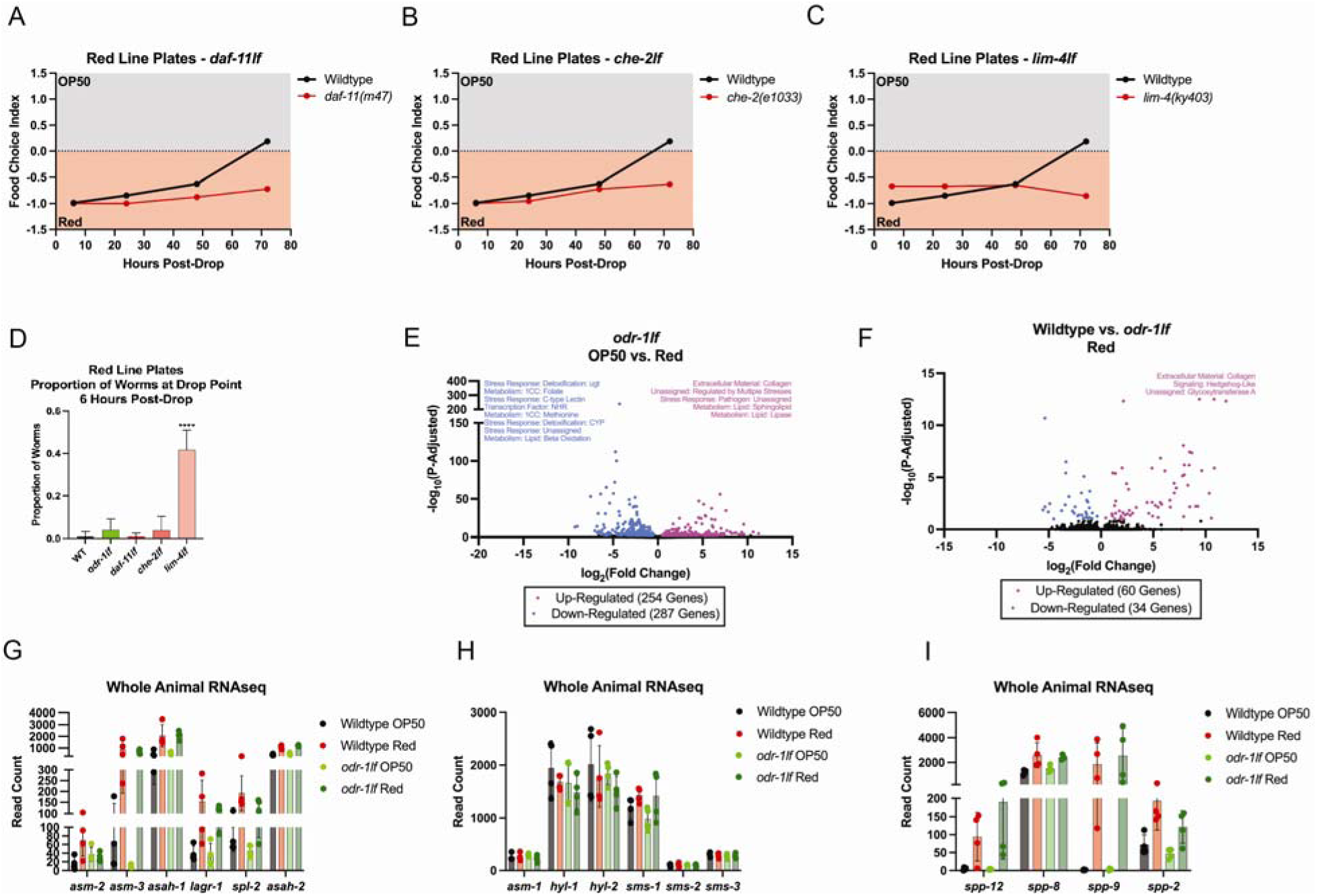
Loss of chemosensation results in changes to food choice and gene expression. (**A-C**) When compared to wildtype worms, *daf-11lf* (**A**), *che-2lf* (**B**), and *lim-4lf* (**C**) do not move away from *Methylobacterium* on Red Line Plates. (**D**) When compared to WT, only *lim-4lf* mutants show a movement defect in food choice assays as measured by the proportion of worms that don’t move from the drop point on the plate after 6 hours. (**E**) *odr-1lf* mutants display a unique transcriptomic signature when fed either *E. coli/*OP50 or *Methylobacterium/*Red. Upregulated pathways are stress response and sphingolipid and lipase metabolism while methionine and beta-oxidation pathways are downregulated. (**F**) There are very few changes between wildtype worms and *odr-1lf* mutants raised on *Methylobacterium*/Red (total of 94 genes differentially changed on the diet), which is highlighted in the whole animal RNAseq read count graphs (**G-I**).

**Figure S7.**
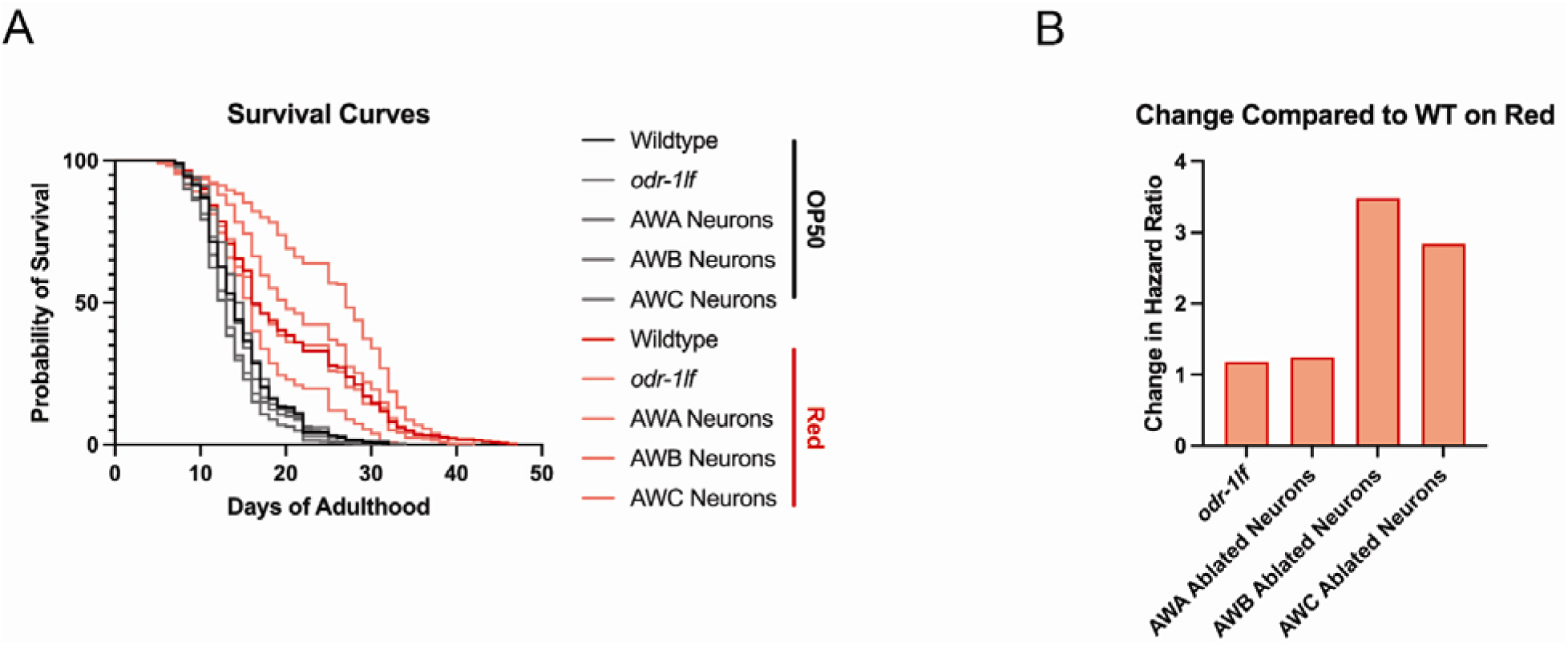
*Methylobacterium* diet-dependent lifespan extension is not dependent upon functional ODR-1, AWA, AWB, or AWC neurons. (**A**) Lifespan of chemosensory mutants are similar to WT when raised on OP50 but change when raised on *Methylobacterium*. (**B**) Change in hazard ratio compared to wildtype worms on *Methylobacterium* reveals AWB-ablated neuron mutants have the largest lifespan extension.

**Table S1.** Lipidomics of WT *C. elegans* fed *E. coli*/OP50 or *Methylobacterium/*Red.

**Table S2.** RNA-Sequencing Analysis of WT and *odr-1lf* mutants on *E. coli*/OP50 and *Methylobacterium/*Red.

**Table S3.** Untargeted Metabolomics in all bacterial genera: *E. coli*/OP50, *E. coli*/HT115, *E. coli*/HB101, *Methylobacterium*/Red, *Xanthomonas*/Orange, *Sphingomonas*/Yellow.

**Table S4.** AWB/AWC-Enriched RNA-Sequencing Analysis of *odr-1p::*RFP on *E. coli*/OP50 and *Methylobacterium/*Red.

**Table S5.** Survival curve comparisons.

**Table S6.** Strains used in this study.

## MATERIALS & METHODS

### *C. elegans* strains and maintenance

*C. elegans* were raised on 6 cm nematode growth media (NGM) plates supplemented with streptomycin and seeded with each bacterial diet. For experiments, nematode growth media plates without streptomycin were seeded with each bacterium at the optical density of 0.8 A_600_. All worm strains were grown at 20°C and unstarved for at least three generations before being used. Strains used in this study are outlined in Table S5 below. Some strains were provided by the CGC, which is funded by NIH Office of Research Infrastructure Programs (P40 OD010440).

*E. coli* strains used: OP50, HT115(DE3), HB101; unless specifically stated otherwise the “*E. coli* diet” refers to the standard OP50 bacteria. Bacteria were isolated from stock plates in the laboratory and selected for with antibiotics before inoculating. Pure monocultures were sequenced to ensure no contamination was present using the 16S primer pair 337F (GACTCCTACGGGAGGCWGCAG) and 805R (GACTACCAGGGTATCTAATC) and identified using the blastn suite on the NCBI website.

### Bacterial Growth & Treatment

Bacterial strains were grown as previously described in Stuhr & Curran 2020 [19]. In brief, *E. coli/OP50* was grown at 37°C overnight in LB supplemented with streptomycin and seeded at an optical density of 0.8 A_600_ onto NGM plates without antibiotics. *Methylobacterium* was grown in LB supplemented with ampicillin for two days at 37°C before being seeded at an optical density of 0.8 A_600_. For experiments where *Methylobacterium* is seeded on food choice plates with other bacterial diets, it is seeded 5 days prior to the other bacteria due to its slow growth on a plate.

For resuspension of *E. coli/OP50* in *Methylobacterium* growth medium, both bacteria were grown as described above. After required inoculation time, bacteria were spun down at 4°C, 4500 rpm for 10 minutes before transferring the supernatant and resuspending OP50 in the *Methylobacterium* growth media.

Killed *Methylobacterium* and *E. coli* treatments were adapted from Stuhr & Curran 2023 [67]. In brief, 800 mL of live bacteria was seeded onto NGM without antibiotics and allowed to grow. Bacteria was then scraped off of the plates and resuspended in sterile M9 before adding 10% PFA for a final concentration of 0.25% PFA. Bacteria was shaken at 37°C in a shaking incubator at 200 rpm for 2 hours. Bacteria was then spun down and resuspended in the same volume of sterile M9 with two subsequent wash steps before final resuspension at 5x concentration in sterile M9.

### Fatty Acid Supplementation

Although multiple methods were tested, for all experiments performed we first seeded plates with bacteria and allowed to grow overnight. 1 hour before experimentation, we supplemented plates with 40mM or 10mM of valeric acid, palmitic acid, salicylic acid, vaccenic acid, oleic acid, or ethanol as the control (all fatty acids were resuspended in ethanol). A total of 30uL was added to the top of the seeded bacteria and dried under a hood before dropping synchronized worms on the plates for food choice analyses.

### Transgenic Animal Generation

To generate tissue-specific expression of WT *odr-1* in the AWB neurons, the coding and 3’ UTR of functional *odr-1* was cloned between the *str-1p* 5’ regulatory sequence and an SL2::WrmScarlet marker and injected along with a *myo-2p::mCherry* marker into *odr-1lf* animals. Pharyngeal expression of mCherry was used to identify transgenic and nontransgenic siblings for food choice testing. Similarly, pan-neuronal expression of functional *egl-3* or gain-of-

function *egl-30* was generated with the use of *rgef-1p* 5’ regulatory sequence in place of the *str- 1p* sequence. *spp-9* injections were performed with a PCR amplified *spp-9p* 5’ regulatory sequence and coinjected with *myo-2p::mCherry* to identify transgenic offspring.

### Movement Measurements – Crawling

Worms were egg prepped and eggs were allowed to hatch overnight for a synchronous L1 population. The next day, worms were dropped onto plates seeded with OP50. Worms were then allowed to grow until each time point (48 h post-drop for L4s, 72 h post-drop for Day 1 Adults, 120 h post-drop for Day 3 adults and 168 h post-drop for Day 5 adults). Once worms were the required stage of development, 30-50 worms were washed off of a plate in 50 uL of M9 with a M9+triton coated P1000 tip and dropped onto an unseeded NGM plate. The M9 was allowed to dissipate, and worms roamed on the unseeded plate for 1 hour before imaging crawling. Crawling was imaged with the MBF Bioscience WormLab microscope and analysis was performed with WormLab version 2022. Worm crawling on the plate was imaged for 1 minute for each condition at 7.5 ms. Worm crawling was analyzed with the software and only worms that moved for at least 90% of the time were included in the analysis.

### Food choice assays

Bacteria was grown overnight in liquid culture of LB with corresponding antibiotics. The next day, bacteria were collected at the log phase, 30uL of each bacterium was seeded onto NGM plates with no antibiotics at 0.8 optical density and allowed to grow overnight. All bacteria were seeded 2 cm from the center point on a 6 cm plate. For the line plates, a glass Pasteur pipette was bent with a flame at a 90-degree angle and then used to transfer bacteria as a line in the center of the plate. The line was 2 cm from the drop point and 2 cm from the spot of food. Once food choice assay plates were seeded and allowed to grow overnight, worms were egg prepped and eggs were allowed to hatch overnight for a synchronous L1 population. The next day, L1s were dropped into the center of the NGM plate and counted. The plate was then checked at 6, 24, 48, and 72 hours to observe the location of worms. If worms were found on the bacterial lawn, then those worms were counted as on that food. Worms found outside bacterial lawns were counted as not on food. The proportion of worms found on each food or off of food was then calculated and graphed. Each assay was done in biological triplicate with technical triplicates for a total of nine plates.

For transgenic strains, food choice was performed slightly differently. Transgenic animals were bleach spotted onto the food choice plate and food choice was examined 48 and 72 hours after L1s hatched. These experiments were conducted under a fluorescent microscope in order to check food choice of the transgenic population on the plate. Nontransgenic siblings were not used in the calculation of food choice indices.

### Lifespan Assays

Worm strains were egg prepped to generate a synchronous L1 population overnight. Worms were kept at 20°C and at the L4 stage, and 50-60 worms were transferred to plates supplemented with 30 uM FUdR to suppress progeny production. Worms were scored daily for survival by gentle prodding with a platinum wire if worms were not freely moving on their own. Animals that burst or crawled to the side of the plate were censored from the analysis. Analysis was done with Long-rank (Mantel-Cox) test and hazard ratios were calculated with Mantel-Haenszel.

### Lipidomics

Lipid extraction and GC/MS of extracted, acid-methanol-derivatized lipids was performed as described previously[68, 69]. Briefly, 5000 synchronous L4 animals were sonicated with a water

bath sonicator on high intensity in a microfuge tube in 250 µL total volume. Following sonication, lipids were extracted in 3:1 methanol:methylene chloride following the addition of acetyl chloride in sealed borosilicate glass tubes, which were then incubated in a 75°C water bath for 1 hr. Derivatized fatty acids and fatty alcohols were neutralized with 7% potassium carbonate, extracted with hexane, and washed with acetonitrile prior to evaporation under nitrogen. Lipids were resuspended in 200 µL of hexane and analyzed on an Agilent GC/MS equipped with a Supelcowax-10 column. Fatty acids and alcohols are indicated as the normalized peak area of the total of derivatized fatty acids and alcohols detected in the sample. Based upon power calculation for pairwise comparison, a minimum n of 3 biological replicates (per group) was chosen to satisfy α=0.05, β=0.2, and effect size = 50% with σ=20%. Analyses were blinded to the investigator conducting the experiment and mass spectrometry calculations until the conclusion of each experiment when aggregate statistics were computed.

### Metabolomics

Metabolomics was conducted on bacterial samples, with 5 replicates per sample. Briefly, bacterial were grown up in 5 1000mL aliquots and spun down into a pellet. Pellets were frozen and sent to Creative Proteomics for untargeted metabolomics via ultimate 3000LC combined with Q Exactive MS from ThermoScientific for UPLC-MS analysis. When samples arrived at the facility, they were thawed and vortexed before a 30-minute sonication, 60-minute freeze, and then a 15-minute centrifugation. Finally, 200uL of supernatant was combined with 1mg/mL DL-o-Chlorophenylalanine. Raw data was acquired and aligned using Compound Discover (3.0 Thermo) based on m/z value and retention time. Significant compounds were selected if they had a fold change greater than 2 and a p-value<0.05[70].

### RNAi Treatment

RNAi treatment was performed as previously described [71]. Briefly, HT115 bacteria containing specific double stranded RNA-expression plasmids were seeded on NGM plates containing 5mM isopropyl-β-D-thiogalactoside and 50μgml^-1^ carbenicillin. RNAi was induced at room temperature for 24 h. Synchronized L1 animals were added to those plates to reduce expression of the indicated genes. Worms were raised for one generation on RNAi and then day 1 adults were bleached to the food choice plates to measure food choice index of the progeny from L1 to day 1 adult stage.

### RNAseq Analysis

RNAseq analysis was conducted as outlined in Stuhr & Curran 2020 [19]. Worms were egg prepped and eggs were allowed to hatch overnight for a synchronous L1 population. The next day, L1s were dropped on to seeded NGM plates and allowed to grow 48 hours to the L4 stage before collection. Animals were washed 3 times with M9 buffer and frozen in TRI reagent at - 80°C until use. Animals were homogenized and RNA extraction was performed via the Zymo Direct-zol RNA Miniprep kit (Cat. #R2052). Qubit™ RNA BR Assay Kit was used to determine RNA concentration. The RNA samples were sequenced and read counts were reported by Novogene. Read counts were then used for differential expression (DE) analysis using the R package DESeq2 created using R version 3.5.2. Statistically significant genes were chosen based on the adjust p-values that were calculated with the DESeq2 package. Gene Ontology was analyzed using the most recent version of WormCat 2.0 [72].

AWB- and AWC-enriched RNAseq was performed as previously described [73, 74]. In brief, approximately 250,000 L4 synchronized *odr-1::RFP* worms grown on *E. coli/OP50,* or *Methylobacterium/Red* were washed with M9 6 times to remove residual bacteria. Animals were then pelleted and Cell Isolation Buffer(20mM HEPES, 0.25% SDS 200mM DTT 3% Sucrose pH8) was added to worms. Worms were incubated in Cell Isolation Buffer for 2min. Initial lysis was quenched by washing with M9 6 times. Worm pellet was resuspended in 20mg/ml Pronase and digested for 20 mins with vigorous pipetting every 5mins through a P1000 tip. Pronase digestion was quenched by resuspending in FBS. Cells were pelleted by centrifuging at 550rcf and resuspended in fresh FBS, Cells were passed through a 40-micron cell strainer. DAPI was added to cells to assess viability. Cells were then sorted on a Bio-Rad S3e FACS system. Neurons were homogenized and RNA extraction was performed via the Zymo Direct-zol RNA Miniprep kit (Cat. #R2052). Qubit™ RNA BR Assay Kit was used to determine RNA concentration. Low input RNA libraries were prepped using the Ovation SoLo RNA library kit from Tecan Genomics. The RNA libraries were sequenced by Novogene. Raw paired-end reads were quality trimmed and adapter trimmed using timmomatic v0.39. Quality trimmed reads were aligned to the *C. elegans* reference genome using STAR 2.7.6a. Mapped reads were counted via HTseq v2.0.2 union mode. Read counts were then used for differential expression (DE) analysis using the R package DESeq2 created using R version 3.5.2. Statistically significant genes were chosen based on the adjust p-values that were calculated with the DESeq2 package. Gene Ontology was analyzed using the most recent version of WormCat 2.0.

### Statistics and Reproducibility

Data are presented as mean ± SEM. Comparisons and significance were analyzed in Graphpad Prism 8. Comparisons between more than two groups were done using ANOVA. For multiple comparisons, Tukey’s multiple comparison test was used, and p-values are *p<0.05 **p<0.01 *** p<0.001 ****p<0.0001. Lifespan comparisons were done with Log-rank test. Sample size and replicate number for each experiment can be found in figures and corresponding figure legends. This information is also in the experimental methods. Exact values for graphs found in the main figures can be found in Supplementary Data.

### Author Contributions

**Competing Interests:** All authors declare that they have no competing interests.

**Data and Materials Availability:** All data are available in the main text or the supplementary materials. RNA-sequencing data are deposited into the Gene Expression Omnibus (GEO). All other relevant data is available upon request from the corresponding author.

## Notes

### Competing Interest Statement

The authors have declared no competing interest.

